# Post-Inhibitory Rebound by δ-Cells Converts Inhibition into Excitation and Confers Islet Plasticity

**DOI:** 10.1101/2025.11.10.687745

**Authors:** Alejandro Tamayo-Garcia, Dayleen Hakim-Rodriguez, Elisabeth Pereira, Sergio Camacho-Becerra, Luciana Mateus Gonçalves, Per-Olof Bergreen, Madina Sokolov, Rafael Arrojo e Drigo, Oscar Alcazar, Rayner Rodriguez-Diaz

**Affiliations:** Division of Endocrinology & Metabolism, Department of Medicine, University of Miami Miller School of Medicine, Miami, FL, USA; The Rolf Luft Research Center for Diabetes and Endocrinology, Karolinska Institute, Stockholm, Sweden; Department of Molecular Physiology and Biophysics, Vanderbilt University School of Medicine, Nashville, TN, USA; Diabetes Research Institute, University of Miami Miller School of Medicine, Miami, FL, USA

**Keywords:** δ-cells, somatostatin, post-inhibitory rebound (PIR), paracrine signaling, insulin secretion, glucagon, excitability, metabolic resilience, endocrine plasticity

## Abstract

The coordination of hormone secretion within the pancreatic islet remains incompletely understood. Here we identify post-inhibitory rebound (PIR), a transient excitatory overshoot of glucoregulatory hormones following inhibitory signaling, as a fundamental and previously unrecognized feature of islet physiology. Human islets from donors with type 2 diabetes displayed concomitantly elevated insulin and somatostatin secretion with tightly correlated dynamics, a pattern reproduced in islets from high-fat-diet–fed mice. Using a combination of cell-type-specific chemogenetic and optogenetic approaches, high-resolution triple-hormone perifusion, Ca²⁺ imaging, in vivo islet grafts, and physiological ligands, we demonstrate that transient δ-cell activation or inhibition consistently triggers rebound excitation of β- and α-cells. These responses generate glucose-dependent, synchronized oscillations of insulin and glucagon secretion that are partially mediated by somatostatin receptor signaling. Mechanistic insights obtained in mouse islets were recapitulated using translatable approaches in human islets, establishing the relevance of this dynamic signaling framework across species. In vivo, selective δ-cell modulation bidirectionally regulated systemic glucose tolerance, confirming δ-cell control over endocrine output. These findings support a model in which somatostatin signaling encodes hormone output through temporal dynamics rather than tonic inhibition. By reframing somatostatin as a dynamic regulator of excitability, we identify δ-cell–driven rebound as a mechanism that converts inhibition into coordinated endocrine output. Together, this work establishes a revised framework for intra-islet paracrine crosstalk in which temporal dynamics, rather than static inhibition, govern endocrine coordination.

**Significance Statement:** Hormone secretion in the pancreatic islet has long been explained by inhibitory feedback loops, with somatostatin viewed as a static suppressor of insulin and glucagon release. Our study overturns this paradigm by identifying post-inhibitory rebound (PIR) as a dynamic property of intra-islet signaling. δ-cell–driven PIR synchronizes α- and β-cell activity, enabling adaptive, glucose-dependent hormone release and revealing that somatostatin confers system plasticity when needed. This work establishes a revised model of intra-islet paracrine crosstalk in which δ-cell–driven rebound dynamics, rather than static inhibition, determine endocrine coordination and system plasticity. Our discovery reframes inhibition as a rhythmic and regenerative force, positioning PIR as a potentially generalizable mechanism of excitability across biological systems.

**Highlights:**

- δ-cells convert inhibition into excitation. Transient somatostatin signals trigger post-inhibitory rebound (PIR) in β- and α-cells, generating glucose-dependent synchronized hormone oscillations.
- Provides a mechanistic explanation for a long-standing paradox in human type 2 diabetes. Elevated basal insulin and somatostatin secretion are positively correlated; PIR explains this co-hypersecretion.
- Asymmetric δ-cell control confers islet plasticity. Stronger inhibition of α-cells versus β-cells enables balanced insulin/glucagon output during feeding or glucagon-skewed surges for hypoglycemia protection.
- Multi-scale validation across species. Chemogenetics, optogenetics, native ligands, Ca²⁺ imaging, in vivo grafts, and a data-informed phenomenological model demonstrate conservation from mouse to human islets.
- Inhibition is instructive. A revised model of intra-islet paracrine crosstalk in which cell-type–specific temporal dynamics encode hormone output through rebound.

**Graphical Abstract:** Somatostatin, long considered a static inhibitor of growth and secretion, instead acts as a dynamic modulator that transforms inhibition into robust, context-dependent excitation. In the islet, amplified SST pulses drive rebound excitation to promote balanced insulin and glucagon output during anabolic demand, while dampening produces glucagon-skewed surges for protection against hypoglycemia. Inhibition in the islet is not merely suppressive but instructive, encoding future hormone output through rebound dynamics. The same pulse–rebound principle may operate in SST-rich systems such as the brain, gut, and tumors, where dynamic modulation of SST signaling confers adaptability and robustness rather than tonic suppression.

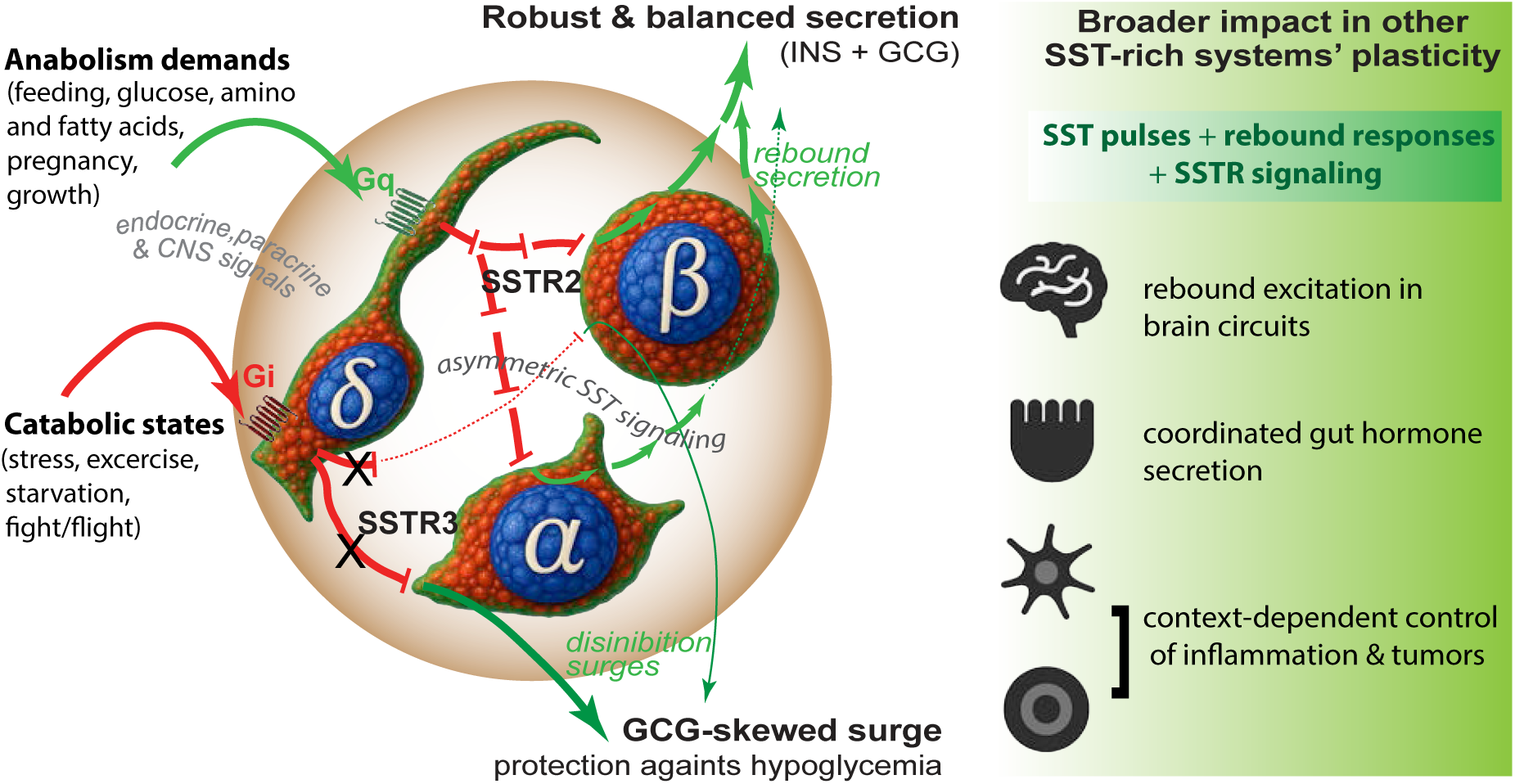

## INTRODUCTION

The maintenance of blood glucose within a narrow physiological range is essential for survival (Röder et al., 2016; Matschinsky & Wilson, 2019). Glucose availability fluctuates with feeding–fasting cycles, activity, and metabolic demand, yet systemic balance is maintained through coordinated regulation by liver, skeletal muscle, adipose tissue, and hypothalamus (Morton et al., 2014; Schwartz et al., 2013). Central to this network is the pancreatic islet of Langerhans, the body’s principal glucose sensor (Rodriguez-Diaz et al., 2018; Noguchi & Huising, 2019; Gao et al., 2021).

Within the islet, α-, β-, and δ-cells form a densely interconnected network integrating hormonal, neural, and metabolic cues to orchestrate glycemia (Hauge-Evans et al., 2009; Benninger & Piston, 2014; Rodriguez-Diaz & Caicedo, 2014). β-cells secrete insulin to promote glucose uptake and storage, while α-cells release glucagon to stimulate hepatic glucose production and as key insulinotropic paracrine signal (Rodriguez-Diaz., et al 2018). Their secretory patterns are pulsatile, a hallmark of efficient endocrine signaling (Hellman, 2009; Edgerton et al., 2021). Disruption of these rhythms is among the earliest abnormalities in type 2 diabetes (Matthews et al., 1983; Matveyenko et al., 2012; Satin et al., 2015). Yet the mechanisms generating and synchronizing these oscillations in vivo remain incompletely understood (Henquin, 2000; Hill & Hill, 2024).

Islet function depends critically on intra-islet communication (Hauge-Evans et al., 2009; Rodriguez-Diaz et al., 2018; Noguchi & Huising, 2019). Each endocrine population acts as both signal generator and receiver, creating a self-regulating microcircuit that maintains the glycemic set point (Pipeleers et al., 1985a,b; Gao et al., 2021). Among these, δ-cells have historically received the least attention. Representing 5–10% of islet cells, they secrete somatostatin (SST), a peptide long considered a static inhibitory brake acting via Gi/o-coupled SSTR2 and SSTR5 receptors to suppress both insulin and glucagon (Strowski & Blake, 2008; Vergari et al., 2019, 2020; Rorsman & Huising, 2018). Yet, as we recently emphasized, such a static framework fails to capture the precision, pulsatility, and dynamic paracrine interactions that define islet function (Tamayo-Garcia et al., 2025).

Accumulating evidence reveals a striking paradox. In both type 2 diabetes (T2D) and type 1 diabetes (T1D), δ-cells exhibit a hypersecretory phenotype, with elevated basal somatostatin output. In T2D this is associated with hyperinsulinemia (this study), whereas in T1D it contributes to impaired counter-regulation during hypoglycemia (Hill et al., 2024; Tegehall et al., 2025). The underlying mechanisms remain incompletely understood and may involve receptor desensitization, altered negative-feedback loops, or other compensatory adaptations. Several lines of evidence have speculated that post-inhibitory rebound (PIR)-like phenomena could contribute to β-cell hyperexcitability in T2D (Nichols & Remedi, 2012). Here we directly test this possibility and demonstrate that somatostatin is not merely a static brake but a dynamic modulator. Using cell-type-specific chemogenetic and optogenetic tools, high-temporal-resolution triple-hormone perifusion, in vivo graft models, Ca²⁺ imaging, native-ligand validation, and mathematical modeling, we show that transient δ-cell-derived inhibitory signals reliably trigger **post-inhibitory rebound (PIR)**—a transient excitatory overshoot in β- and α-cells. PIR converts somatostatin-mediated inhibition into synchronized, glucose-dependent excitation, generating adaptive hormone oscillations that confer plasticity to the islet.

By establishing that somatostatin, through dynamic secretion patterns, post-inhibitory rebound, disinhibition, and differential receptor sensitivity, can generate a broad spectrum of context-dependent outcomes, we reveal it as a powerful modulator that confers significant plasticity and adaptability to the islet and, potentially, to many other SST-regulated systems where it has long been viewed solely as a potent inhibitor. In this framework, inhibition is not a terminal state, but a preparatory phase that encodes future excitation.

## RESULTS

### Paradoxical co-elevation of basal insulin and somatostatin secretion in human type 2 diabetes islets

Type 2 diabetic human islets exhibit elevated basal insulin and somatostatin secretion with impaired glucose responsiveness, revealing an unexpected positive coupling despite the canonical inhibitory action of somatostatin.

The pancreatic δ-cell has classically been viewed as an inhibitory node within the islet of Langerhans, exerting paracrine suppression of both α- and β-cells through secretion of somatostatin, a potent inhibitory peptide acting primarily via SSTR2 and SSTR5 receptors (Strowski & Blake, 2008; Vergari et al., 2019; 2020). By constraining insulin and glucagon release, δ-cells help fine-tune islet hormone ratios and preserve metabolic balance. Given that disordered secretion of both insulin and glucagon contributes to type 2 diabetes (T2D) pathophysiology, we investigated whether δ-cell activity and somatostatin secretion are altered under diabetic conditions.

To address this, we performed high-resolution perifusion experiments in human islets isolated from non-diabetic (ND) lean donors and from individuals with T2D (**Figure 1**, **S1 and Table S1**). Insulin and somatostatin were measured simultaneously across a glucose ramp (3 → 11 mM) to capture both basal and glucose-stimulated secretion (**Fig. 1A–B**). In ND islets, hormone release displayed the expected dynamic profile: low basal secretion with robust glucose-stimulated insulin secretion (GSIS) and a more modest somatostatin response (**Fig. 1C**–**C1**). By contrast, T2D islets displayed a striking phenotype: both basal and glucose-stimulated secretion of insulin and somatostatin were markedly elevated compared with ND islets (**Fig. 1D–D1**). Importantly, while ND islets showed no correlation between basal insulin and somatostatin release, secretion in T2D was tightly coupled (**Fig. 1E**; r² = 0.42, p = 0.0035), indicating coordinated hyperactivity of β- and δ-cells and a functional coupling rather than classical paracrine inhibition. Despite this hypersecretion, total islet content of insulin and somatostatin was reduced in T2D, whereas glucagon content remained unchanged (**Fig. 1G–G2**). Area-under-the-curve analyses further confirmed a surplus of absolute hormone secretion levels in T2D that contrasted with the impaired responsiveness (**Fig. 1F–F1**; **p<0.01**). These findings indicate a state of β- and δ-cell hyperexcitability relative to α-cells, consistent with a shift in intra-islet functional hierarchy, in which elevated output is accompanied by depleted hormone stores.

**Figure 1.**
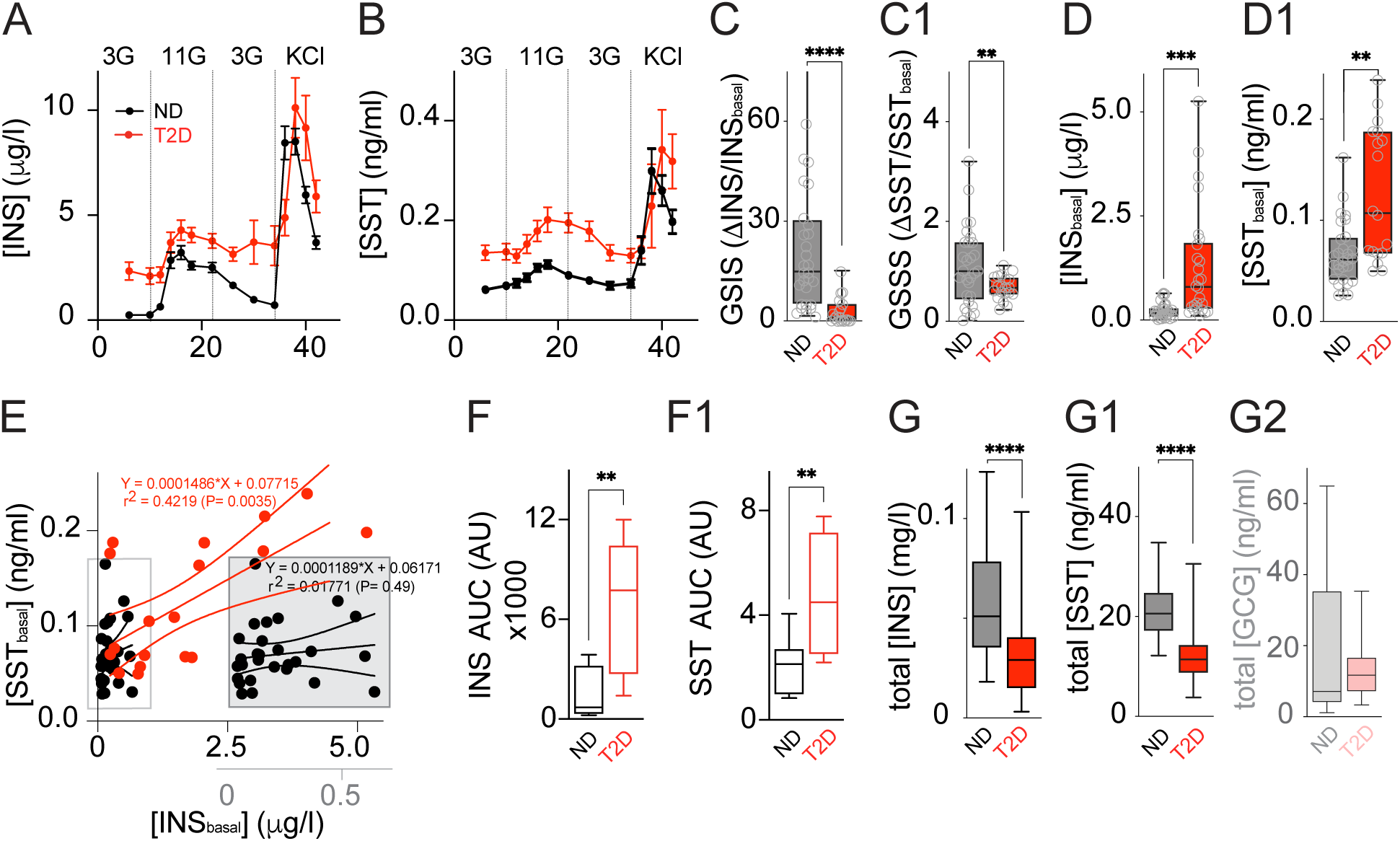
Type 2 diabetic human islets exhibit elevated basal insulin and somatostatin secretion with impaired glucose responsiveness, revealing a paradox resolved by δ-cell dynamics. **(A–B)** Perifusion profiles of insulin and somatostatin secretion in non-diabetic (ND, black, n=15) and type 2 diabetic (T2D, red, n=15) human islets during sequential 3 mM, 11 mM, 3 mM glucose and KCl stimulation (mean ± SEM). **(C–C1)** Glucose-stimulated insulin secretion (GSIS) and somatostatin secretion (GSSS) indices (stimulated/basal ratio). T2D islets show markedly impaired responses (****p<0.0001, **p<0.01, unpaired t-test). **(D–D1)** Basal insulin and somatostatin levels at 3 mM glucose are significantly elevated in T2D islets (***p<0.001, **p<0.01). **(E)** Positive correlation between basal insulin and somatostatin in T2D islets (r² = 0.42, p = 0.0035), absent in ND islets. **(F–F1)** Area under the curve (AUC) for total insulin and somatostatin secretion during glucose ramps (**p<0.01), indicating a surplus of absolute hormone secretion levels in T2D that contrasts with the impaired responsiveness shown above. **(G–G2)** Post-perifusion islet hormone content of 150 IEQ (adjusted by variation in protein content for each preparation): reduced insulin and glucagon, but increased somatostatin in T2D (*p<0.05).

This hypersecretory trend was recapitulated in murine models. Islets and pancreatic slices from high-fat diet (HFD)-fed mice showed elevated basal secretion of both insulin and somatostatin compared with regular chow-fed (RD) controls (**Figure S2**).

This unexpected coexistence of elevated basal insulin and somatostatin in T2D presents a paradox under the classical view of somatostatin as a tonic inhibitor of islet hormone secretion, suggesting that δ-cell signaling operates through fundamentally different dynamics in the diabetic state.

These observations raise a fundamental question: how can a classically inhibitory δ-cell signal remain positively associated with β-cell output? We reasoned that static models of paracrine inhibition may not capture the dynamic nature of intraislet signaling, particularly under conditions of altered excitability. In addition, prior approaches relying on pharmacological agonists may have been limited by off-target or non-cell-specific effects, potentially obscuring cell-resolved interactions. To directly interrogate the temporal and cell-specific mechanisms underlying δ-cell–mediated regulation, we employed a chemogenetic approach to selectively manipulate α-, β-, and δ-cell activity and resolve their functional interplay in real time.

### Chemogenetic mapping reveals post-inhibitory rebound as a key mechanism of intraislet paracrine crosstalk

To dissect how δ-cells modulate islet communication, we generated transgenic mouse lines expressing the excitatory DREADD hM3Dq selectively in β-cells (INS-Cre), α-cells (GCG-Cre), or δ-cells (SST-Cre). Three F1 crosses were established: INS-DREADD Gq, GCG-DREADD Gq, and SST-DREADD Gq. To complement activation studies, we also generated δ-cell–specific inhibitory the DREADD hM4Di line SST-DREADD Gi to enable bidirectional chemogenetic control of δ-cell activity. Immunocytochemistry confirmed cell-specific plasma-membrane localization of HA-tagged hM3Dq (**Fig. 2A, E, I**).

**Figure 2.**
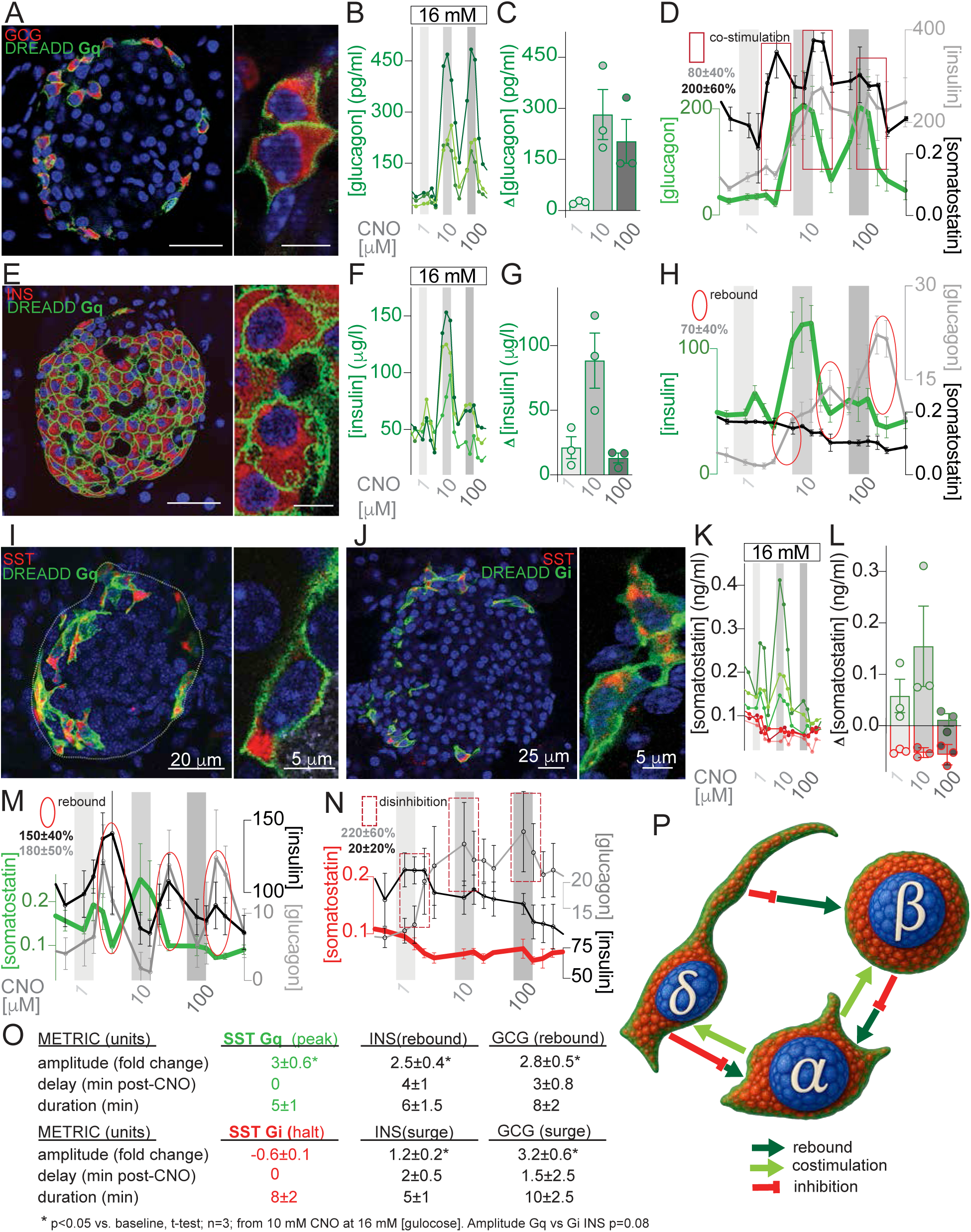
Comprehensive chemogenetic mapping of α-, β-, and δ-cell interplay reveals post-inhibitory rebound as a novel mechanism of intraislet paracrine crosstalk, adding nuance into intraislet regulation. Experiments validate cell-type-specific DREADD expression and homotypic secretion responses, while examining secondary induced responses in neighboring cells (n=3-5 per condition). Dynamic secretion of insulin, somatostatin, and glucagon was measured in parallel via perifusion at 1 mM, 3 mM, and 16 mM glucose, with results shown here for 16 mM where effects are most prominent. Clozapinbe-N-Oxide (CNO) was applied in 4 min pulses with titration followed by washout. **(A–D)** Validation of cell-type specific expression of GCG-DREADD Gq in α-cells by ICC (A) and time courses of glucagon secretion upon acute CNO application (B,C). α-cell activation triggers co-stimulatory responses in β- and δ- cells (D) **(E–H)** Validation of cell-type specific expression of INS-DREADD Gq in β-cells (E) and time courses of insulin secretion upon acute CNO application, showing additional surplus/oversoot (F,G). β-cell activation also triggers post-inhibitory rebound (PIR) in α-cells, stimulating glucagon release (H), thought PIR is more prominently observed and further characterized with δ-cell modulation below as the inhibitor par excellence. **(I–L)** Validation of cell-type specific expression of SST-DREADD Gq and SST-DREADD Gi in δ-cells (I,J) and time courses of somatostatin secretion upon acute CNO application (Gq green, Gi red; K,L). **(M–N)** Rebound response of insulin (black) and glucagon (gay) after Gq activation (M) and disinhibition surge (glucagon-skewed) after Gi activation in δ-cells (N). **(O)** Summary of key metrics (amplitude, delay, duration) for SST Gq and Gi effects (mean ± SEM, n=3). **(P)** Novel emerging schematic model of intraislet paracrine crosstalk elicited by chemogenetic activation of α-, β-, and δ-cells together with selective δ-cell inhibition. In addition to well-established co-stimulatory responses, the diagram reveals post-inhibitory rebound excitation (PIR) following Gq activation and differential inhibition (evidenced by disinhibition surges after Gi activation), highlining the asymmetric influence of δ-cells on α- and β-cells. These findings identify PIR as a previously underappreciated mechanism (observed also from β- to α-cells) and support a revised model for the orchestration hormone secretion (detailed in Fig. 3), which forms the experimental basis for the mathematical model of SST pulse-rebound and disinhibition surge dynamics in Fig. 7A–C and the Graphical Abstract.

In perifusion assays, CNO elicited robust dose-dependent increases in homotypic hormone secretion (insulin, glucagon, or somatostatin; maximal at 10 μM, partial desensitization at 100 μM; **Fig. 2B–C, F–G, K–L** and **Figure S3**). Responses were further potentiated under high-glucose conditions; notably, α-cells displayed strong glucagon secretion despite 16 mM glucose (**Fig. 2B–C** and **S3**).

CNO-driven activation produced immediate homotypic responses and delayed heterologous responses across cell types, including both co-stimulatory effects during activation and delayed responses following stimulus withdrawal. α-cell stimulation boosted insulin and somatostatin release, consistent with established paracrine interactions (**Fig. 2D and S3**). δ-cell Gq activation initially suppressed insulin and glucagon secretion, followed by pronounced post-inhibitory rebound (PIR) surges that resulted in net increases in hormone output (**Fig. 2M, O and S3**). β-cell activation similarly induced PIR in α-cells, triggering glucagon release shortly after stimulation (**Fig. 2H and S3**).

Reproducibility was confirmed in two additional independent SST-DREADD Gq lines (**Figure S4**), where CNO consistently evoked acute somatostatin release followed by delayed insulin and glucagon rebounds, demonstrating that these dynamics are intrinsic to δ-cell–mediated inhibition.

δ-cell Gq activation triggered robust somatostatin release followed by pronounced PIR in insulin (2.5 ± 0.4-fold) and glucagon (2.8 ± 0.5-fold; Fig. 2M, O). Conversely, δ-cell Gi inhibition halted somatostatin secretion **(Fig. 2K, L)** and elicited disinhibition surges in insulin (1.2 ± 0.2-fold) and glucagon (3.2 ± 0.6-fold; **Fig. 2N, O**). These asymmetric responses establish a conceptual model of intra-islet paracrine crosstalk in which transient inhibitory signals encode hormone output through pulse–rebound dynamics rather than tonic suppression, as illustrated in **Fig. 2P**, highlighting the asymmetric influence of δ-cells on α- and β-cells.

### δ-cell Gq activation drives post-inhibitory rebound while Gi activation produces disinhibition surges that reveal asymmetric control of hormone output

Building on the comprehensive chemogenetic mapping in **Figure 2**, we next quantified the differential impact of δ-cell Gq and Gi manipulation on islet hormone dynamics at 16 mM glucose (**Figure 3**). Gq activation produced rapid SST peaks (**Fig. 3 A–A1,** green) followed by robust rebound increases in both insulin and glucagon (**Fig. 3 B–B1 and C- C1**, green). In contrast, Gi activation caused a sharp SST halt (**Fig. 3 A–A1,** red) followed by disinhibition surges that were markedly stronger for glucagon than insulin (**Fig. 3 B–B1 and <u>C- C1</u>**<u>, red</u>). To highlight surplus secretion relative to control, individual traces were subtracted from the average control trace, with mean ± SEM shown in pairs (left: individual curves; right: surplus traces). Quantitative metrics confirmed the asymmetry: Gq induced significant rebound (net gain +150–220%), while Gi induced strong disinhibition surges (net gain +300–400% for glucagon) (**Fig. 3D–F**, *p<0.05, t-test). Violin plots illustrated the temporal differences: Gq biphasic (initial inhibition depth after CNO followed by delayed rebounds), Gi concomitant surges (**Fig. 3F–F1**, shadowed gray area represents CNO pulse, *p<0.05 vs baseline, t-test). Panel G schematizes hormone traces following δ-cell manipulation (see **Fig. S5–S6**), assuming sequential CNO pulses modulate endogenous δ-cell pulsatility. These simplified traces define the model hypothesis, illustrating depth, rebound, and surge dynamics recapitulated in **Fig. 7B**.

**Figure 3.**
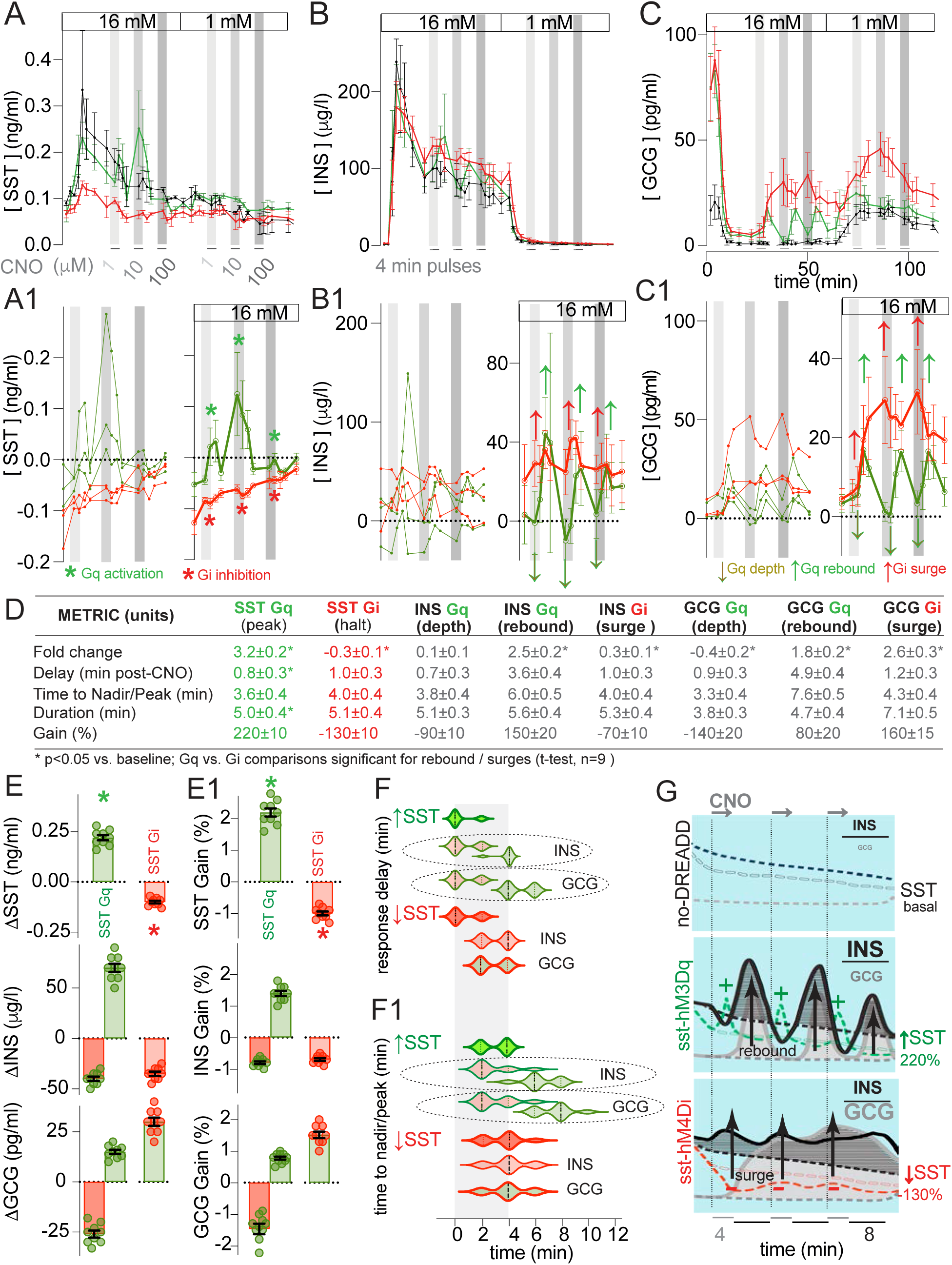
δ-cell Gq activation drives post-inhibitory rebound while Gi activation produces disinhibition surges, revealing asymmetry that underpins balanced versus skewed hormone output. (A–C) Perifusion protocol at 16 mM and 1 mM glucose with acute CNO pulses. Time courses of somatostatin, insulin, and glucagon secretion upon CNO 4 min pulses and 8 min of washout (Gq green, Gi red, control black). **(A1–C1)** Control-subtracted individual traces (left) and mean ± SEM surplus traces (right) of somatostatin, insulin, and glucagon secretion upon CNO (Gq green, Gi red), highlighting Gq depth and rebound, and Gi surge. **(D)** Summary of key metrics (fold change, delay, time to nadir/peak, duration, and net gain) for Gq and Gi effects (mean ± SEM, n=9). Gq activation produces a rapid SST peak with concomitant transient inhibition (depth) followed by delayed (3-8 min) rebound increases in both insulin and glucagon, whereas Gi activation causes a sharp SST halt concomitant with disinhibition surges that are markedly stronger for glucagon than insulin (*p<0.05 vs baseline; Gq vs Gi comparisons significant for rebound/surges, t-test). **(E–E1)** Quantification of net change in amplitude of responses and gains for somatostatin, insulin, and glucagon under Gq and Gi activation. **(F–F1)** Violin plots showing timing of depth, rebounds, or surges following CNO (shadowed gray area represents the CNO pulse). Gq induces biphasic responses with initial inhibition depth right after CNO followed by delayed rebounds (significant delays as per table D), while Gi induces concomitant surges without biphasic pattern (*p<0.05 vs baseline, t-test). **(G)** Manual graphical traces providing a visual impression of the hypothesis for the mathematical model (Fig. 7B): no-DREADD control vs Gq rebound (balanced INS/GCG) vs Gi surge (GCG-skewed). These asymmetric responses (stronger inhibition of α-cells than β-cells) form the experimental basis for the mathematical model of SST pulse-rebound dynamics shown in Fig. 7A–C.

These asymmetric responses, characterized by stronger inhibition of α-cells than β-cells, provided the experimental foundation for the mathematical/phenomenological model of δ-cell pulse-rebound dynamics (**Figure 7**) and directly explain the balanced rebound under Gq and glucagon-skewed surge under Gi observed in vivo.

### Glucose-dependent post-inhibitory rebound of insulin secretion following transient δ-cell activation

To probe the physiological basis of post-inhibitory rebound (PIR) of insulin secretion, we assessed whether this phenomenon is glucose-dependent and mediated by somatostatin (SST). Perifusion of islets from sst-Cre; hM3Dq(Gq) transgenic mice at varying glucose concentrations (3–16 mM) following δ-cell activation with CNO revealed that rebound insulin secretion was absent at basal glucose (3 mM) but robustly emerged under permissive conditions (7–16 mM), with peak amplitudes comparable to first-phase glucose-stimulated insulin secretion (GSIS) or KCl-induced depolarization (**Figure 4A, B**). These findings indicate that PIR operates preferentially under high-glucose, nutrient-permissive conditions, aligning with physiological states requiring enhanced anabolic hormone output.

**Figure 4.**
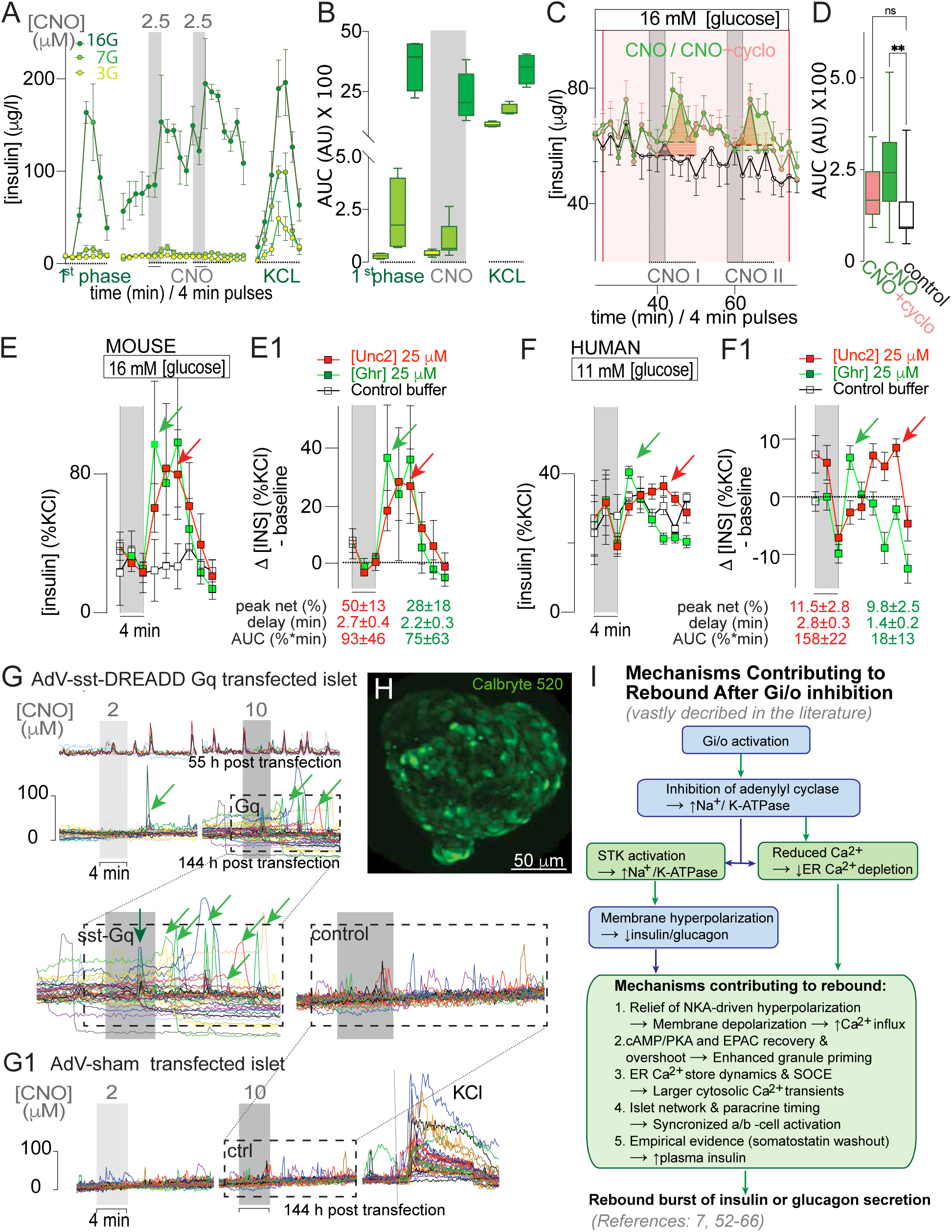
Glucose-dependent post-inhibitory rebound (PIR) of insulin secretion upon transient δ-cell Gq-DREADD activation in transgenic mouse islets is partially mediated by somatostatin; native δ-cell ligands urocortin-3 and ghrelin reproduce PIR with species-specific kinetics in mouse and human islets. (A–B) Perifusion insulin secretion in sst-hM3Dq(Gq) transgenic mouse islets (as in Figs 2–3; n = 3 independent isolations). Traces (A) during initial glucose first-phase stimulation followed by repeated 4-min CNO pulses (2.5 μM) on three different glucose backgrounds (16G, 7G, 3G) and final KCl depolarization. AUC quantification (B) demonstrates clear glucose dependence of the insulin PIR that follows each CNO washout. **(C–D)** On a 16 mM glucose background, CNO pulses alone (red trace) versus CNO plus cyclosomatostatin (SST receptor antagonist, orange shading). Traces (C) and AUC (D) reveal that endogenous somatostatin signaling contributes partially to the insulin PIR (one-way ANOVA/Tukey; **p < 0.01 for CNO vs control; ns for CNO vs CNO + cyclosomatostatin). **(E–F1)** Native δ-cell ligands urocortin-3 (Unc3, 25 μM mouse / 12.5 μM human, red) or ghrelin (Ghr, 25/12.5 μM, green) versus control buffer in mouse (E/E1; n = 3) and human (F/F1; n = 5–6 perifusions from 2 donors) islets on 16 mM or 11 mM glucose. Raw traces (E/F) and control-subtracted Δ[%KCl baseline] (E1/F1). Green and red arrows mark PIR peaks. Insets report PIR metrics (mean ± SEM): peak net gain (%), delay from pulse washout (min), and integrated AUC (%·min over the 6–18 min window post-pulse). In human islets Unc3 produces a markedly larger and more sustained net AUC than ghrelin (***p < 0.001, unpaired t-test). **(G–G1)** Bulk Ca²⁺ imaging (Calbryte 520 dye, 7 mM glucose) in human islets transfected with AdV-Sst-DREADD-Gq (G; 60 h and 160 h post-transfection) or sham vector (G1). Green arrows indicate direct CNO-evoked Ca²⁺ rises; red arrows indicate stereotypical post-inhibitory rebound (PIR) occurring after stimulus withdrawal. Sham-transfected islets showed Ca²⁺ responses that were not dynamically associated with the CNO pulses but efficiently responded to KCl depolarization (viability control). Note that insulin (and glucagon) secretion during perifusion (**Fig. S9**) showed only a non-significant trend toward PIR in these DREADD-Gq-transduced human islets, likely due to incomplete transfection efficiency and limited viral penetration. Validation of δ-cell-specific transduction is shown in Supplementary Figure S9. **(H)** Maximal z-projection of a representative human islet loaded with Calbryte 520 (scale bar, 50 μm). Representative full Ca²⁺ dynamics, quantification, and display are shown in Suppl. FigureS8 and Suppl. Movie 1. **(I)** Mechanistic model (supported by extensive literature) of the subcellular pathways downstream of transient Gi/o signaling that convert inhibition into rebound excitation of insulin secretion (References: 7, 52–66). No new experiments were performed in this panel; the diagram illustrates known mechanisms that can account for the overlooked PIR responses of insulin secretion reported in this study.

Pharmacological interrogation with cyclosomatostatin, a SST receptor antagonist, partially blunted the insulin rebound post-CNO, supporting a contributory role for SST in PIR while indicating the involvement of additional paracrine or intrinsic mechanisms (**Figure 4C, D**; see also **Figure S7** for extended controls including male vs. female donor comparisons and broader pharmacological validation).

### Native δ-cell ligands reveal conserved post-inhibitory rebound mechanisms in mouse and human islets

To validate physiological relevance, we tested native δ-cell–modulating ligands with distinct receptor coupling mechanisms, including urocortin-3 (Ucn3) and ghrelin (Ghr). Both ligands stimulate δ-cell activity and somatostatin release, which in turn engages Gi/o-coupled SSTR signaling in α- and β-cells. In wild-type mouse islets, both ligands elicited clear PIR of insulin secretion with comparable kinetics (**Fig. 4E–E1**).

We next extended these findings to human islets from non-diabetic donors. Ucn3 and Ghr elicited rebound insulin secretion, albeit with markedly divergent kinetics: ghrelin produced a rapid transient peak followed by strong inhibition (net AUC 18 ± 13 %·min, n.s. vs 0), whereas urocortin-3 triggered a slower-onset, sustained excitatory rebound (net AUC 158 ± 22 %·min, ***p < 0.001 vs ghrelin and vs 0; **Fig. 4F–F1**).

To probe the underlying signaling, we performed adenoviral transduction of δ-cells with AdV-Sst-DREADD-Gq (MOI = 50) using an Ad5/F35 chimeric backbone selected to enhance transduction efficiency in primary human islets, which exhibit low coxsackievirus and adenovirus receptor (CAR) expression. δ-cell-specific transduction efficiency, specificity, and functional validation of the responses are shown in **Figure S8**. Bulk Ca²⁺ imaging revealed stereotypical direct responses during CNO application and clear PIR-like transients after stimulus withdrawal (**Figure 4G–H; Suppl. Movie 1**). Insulin secretion showed only a non-significant trend toward PIR (n = 9 islets from 3 independent non-diabetic donors; likely due to incomplete transfection efficiency and limited viral penetration). However, responses were consistently observed across independent donor preparations despite variability in transduction efficiency, indicating robust δ-cell–dependent dynamics that align with mouse data and support conservation of PIR mechanisms across species.

Finally, a schematic integrates established Gi/o signaling cascades that can account for PIR: transient inhibition lowers cAMP/PKA/EPAC, hyperpolarizes β-cells via enhanced Na⁺/K⁺-ATPase activity, suppresses Ca²⁺ influx, and primes for rebound excitation upon relief, yielding amplified insulin release (**Figure 4I**; References 7, 52–66). Collectively, these findings underscore PIR’s role in fine-tuning insulin output under physiological demands and highlight its overlooked impact on islet secretory machinery.

### Orthogonal optogenetic activation of δ-cells elicits calcium rebound and confirms δ-mediated PIR in vivo

To orthogonally validate the post-inhibitory rebound (PIR) mechanism established in vitro (**Figures 3–4**), we transplanted islets expressing channelrhodopsin-2 selectively in δ-cells (or δ-ablated controls) into the anterior chamber of the eye of STZ-diabetic nude mice (**Fig. 5A**). Vascularization and δ-cell specificity were confirmed in intact versus δ-ablated grafts (**Fig. 5B**). Blood glucose time course after transplantation confirmed engraftment restored normoglycemia in both cohorts (**Figure 5C**; endpoint **p<0.001**).

**Figure 5.**
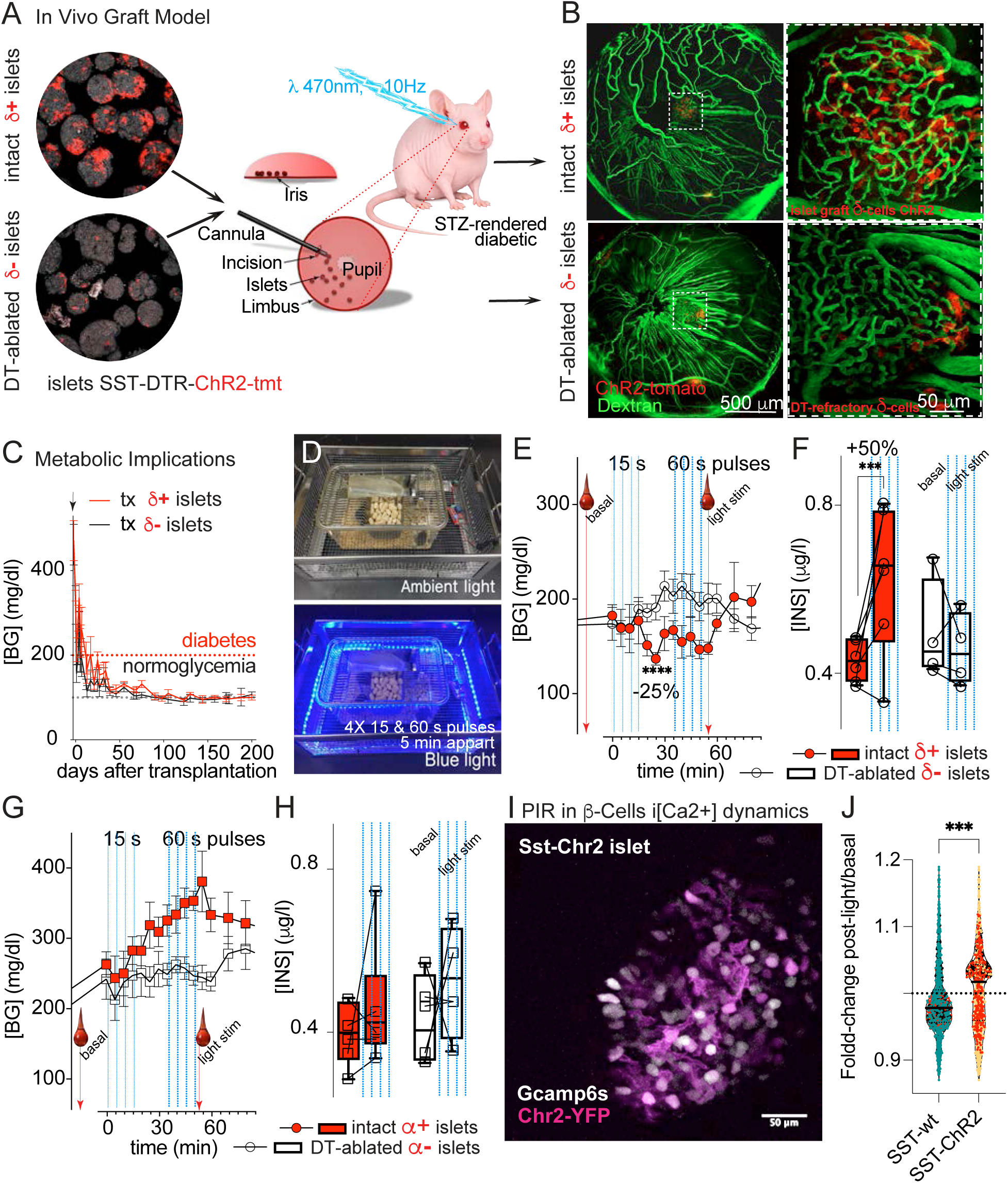
Optogenetic activation of δ-cells in vivo triggers post-inhibitory rebound in β-cells, driving Ca²⁺-dependent insulin secretion and systemic hypoglycemia. Using optogenetics in the anterior chamber of the eye (ACE) graft model to orthogonally validate PIR and its intracellular Ca²⁺ basis under physiological conditions. **(A)** Schematic of the in vivo eye graft model for optogenetic activation of δ-cells: islets from double transgenic mice expressing nChR2-TdTomato and DTR in δ-cells (intact δ-cells or DT-ablated) were transplanted into the ACE of STZ-diabetic nude mice. **(B)** Vascularization and δ-cell specificity in intact (top) vs δ-ablated (bottom) in few islet grafts in contralateral eye used for imaging (ChR2-tdTomato in red, dextran in green). **(C)** Blood glucose time course after transplantation confirms engraftment restores normoglycemia in both cohorts (mean ± SEM, n=4; endpoint **p<0.001). **(D)** Blue-light stimulation setup and protocol (4×15 s + 4×60 s pulses). **(E–F)** Light stimulation induces hypoglycemia (E; Mean ± SEM blood glucose traces, ****p<0.0001 for AUC) and elevated plasma insulin (F; ***p<0.001) exclusively in intact-δ grafts (red) versus δ-ablated (clear). **(G-F)** Specificity control with Gcg-ChR2 grafts (increased blood glucose only in intact α-cell **(I–J)** In vivo GCaMP6s- β-cell Ca²⁺ imaging in SST-ChR2 islets shows enhanced PIR-like Ca2+ peak amplitude fold-change (stimulated/basal) after δ-cell photoactivation (second pulse train further amplifies; n=1,357 control and 1,123 ChR2 cells across n=3 mice/group; ***p <0.001, unpaired t-test with Welch’s; see Suppl. Movie 2).

Blue-light stimulation (4×15 s + 4×60 s pulses; **Fig. 5D**) induced hypoglycemia (**Fig. 5E**; ****p < 0.0001 for AUC) and elevated plasma insulin (**Fig. 5F**; ***p < 0.001) exclusively in intact-δ grafts (red) versus δ-ablated (clear). Specificity was confirmed using Gcg-ChR2 grafts, which increased blood glucose only in intact α-cell grafts (**Fig. 5G–H**).

In vivo GCaMP6s-β-cell Ca²⁺ imaging in SST-ChR2 islets showed enhanced PIR-like Ca²⁺ peak amplitude fold-change (stimulated/basal) after δ-cell photoactivation; the second pulse train further amplified the response (n=1,357 control and 1,123 ChR2 cells across n=3 mice/group; ***p <0.001, unpaired t-test with Welch’s; **Fig. 5I–J**; **Suppl. Movie 2**). These data were obtained by re-analysis of raw recordings from a prior collaborative study (Arrojo e Drigo et al., Nat Commun 10, 3700, 2019), in which the initial short transient inhibition of β-cell Ca²⁺ activity immediately after δ-cell activation was reported. Longer-term analysis now reveals a net PIR-like increase in β-cell Ca²⁺ activity, consistent with the chemogenetic and ligand-induced PIR observed in vitro. This previously unappreciated net excitatory effect explains the unexpected hypoglycemia and elevated plasma insulin we observed during early optogenetic experiments in the eye model.

### In vivo δ-cell manipulation bidirectionally controls insulin secretion and glucose homeostasis

Building on the optogenetic validation of PIR (**Figure 5**), we transplanted islets expressing δ-DREADD-Gq, δ-DREADD-Gi, or control from male and female mouse donors into the anterior chamber of the eye of STZ-diabetic nude mice. All groups restored normoglycemia with comparable baseline glucose and hormone levels both in fasting and during feeding stages (**Fig. 6A–A1, B–E**). As no major sex-specific differences were observed (see **Fig. S9**), results are presented combined. Immunocytochemistry confirmed stable δ-cell identity and membrane-localized HA-DREADD expression months after engraftment (**Figure 6F–F3b**).

**Figure 6.**
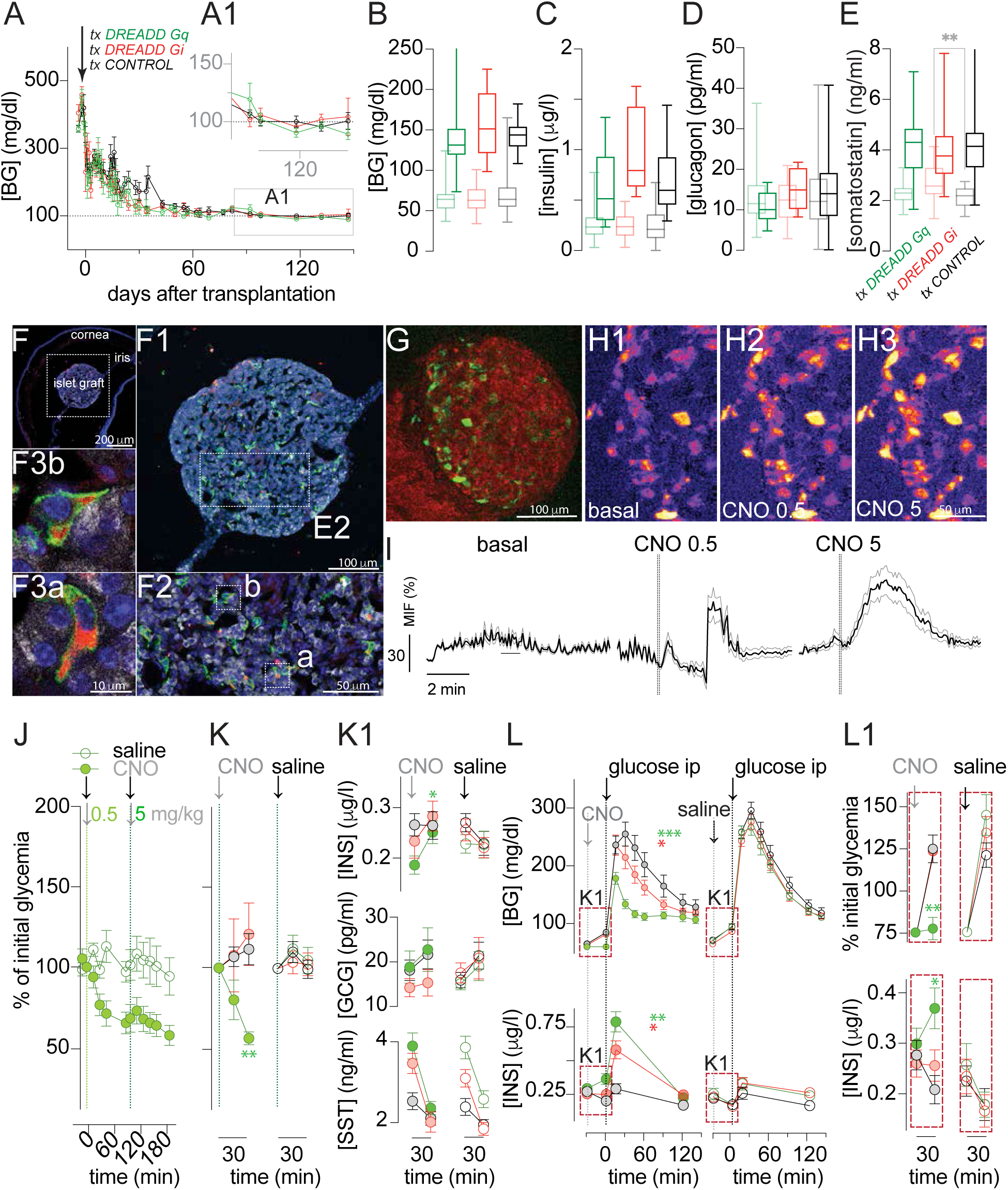
Acute chemogenetic manipulation of δ-cells in vivo reveals PIR-mediated enhancement of islet hormone output and glucose homeostasis. Extending the optogenetic validation of PIR (Figure 5), this figure tests the acute effects of bidirectional δ-cell manipulation using DREADDs in anterior chamber of the eye grafts. **(A–A1)** Longitudinal non-fasting blood glucose after transplantation of δ-DREADD-Gq (green), δ-DREADD-Gi (red), or control (black) islets into STZ-diabetic nude mice shows equivalent restoration of normoglycemia (n≥7/group; consolidated male/female donors). **(B–E)** Fasting (light) and prandial (dark) plasma glucose, insulin, glucagon, and somatostatin levels. **(F–F3b)** Immunocytochemistry of intraocular grafts confirming stable δ-cell identity and membrane-localized HA-DREADD expression months after long engraftment. **(G–I)** In vivo GCaMP3 imaging of δ-DREADD-Gq cells demonstrates dose-dependent Ca²⁺ responses to CNO (0.5 and 5 mg/kg), confirming on-target activation by systemic CNO (G1–G3: representative images; H: mean ΔF/F trace; see Suppl. Movie 3). **(J–K1)** Acute effects of CNO versus saline on glycemia and plasma hormones (insulin, glucagon, somatostatin) in non-fasting conditions. **(L–L1)** Intraperitoneal glucose tolerance test (IPGTT) performed after acute CNO or saline shows improved glucose clearance and enhanced insulin secretion selectively in δ-Gq grafts (*p<0.05, **p<0.01).

In vivo Ca²⁺ imaging of GCaMP3-expressing δ-cells verified that systemic (i.p.) CNO elicits rapid, dose-dependent activation selectively in DREADD-expressing grafts (**Figure 6G–I**; **Suppl. Movie 3**). Acute CNO administration in non-fasting states produced hypoglycemia accompanied by increased insulin secretion in Gq (green) ocular grafts (**Figure 6J–K1**), whereas Gi (red) inhibition had minimal effects. During intraperitoneal glucose tolerance tests, δ-cell Gq activation improved glucose clearance and enhanced insulin secretion relative to control and Gi grafts (**Figure 6L–L1 and S10**; *p<0.05, **p<0.01).

Together, the optogenetic validation in Figure 5 and these acute chemogenetic results demonstrate that δ-cell activation drives PIR-dependent enhancement of islet hormone output and glucose homeostasis in vivo, providing direct physiological support for a dynamic, rebound-based model of intraislet regulation (**Figure 7)**.

**Figure 7.**
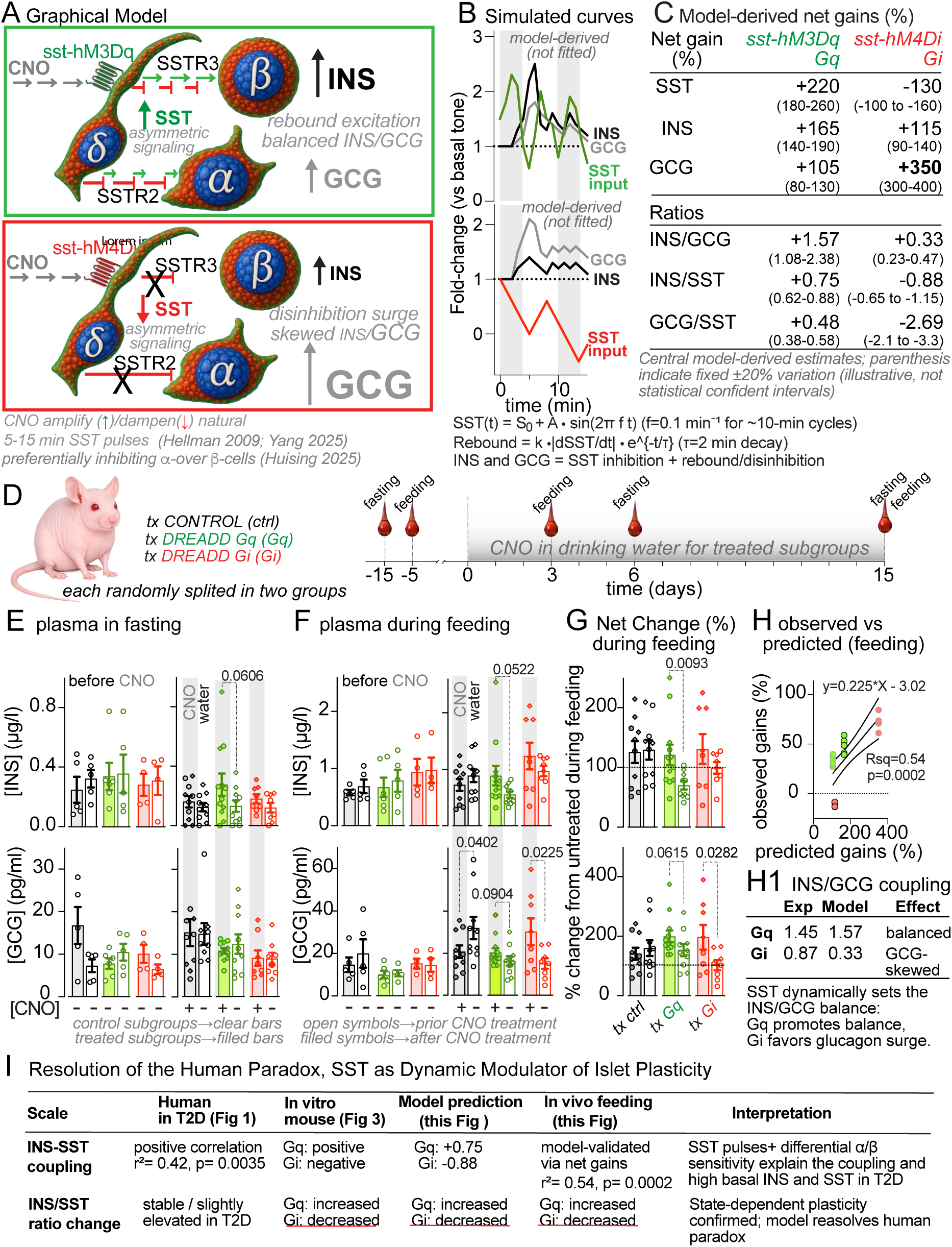
A revised model of intra-islet paracrine crosstalk driven by δ-cell post-inhibitory rebound dynamics which transforms tonic inhibition into dynamic excitation, redefining islet plasticity. **(A)** Graphical model of δ-cell chemogenetic manipulation. Data from Figure 3 showed that SST amplification (Gq) drives rebound excitation with balanced insulin/glucagon coupling favoring insulin, whereas SST dampening (Gi) produces disinhibition surges skewed toward glucagon. SST is secreted in natural pulsatile bursts (5–15 min cycles; Hellman et al., 2009; Yang et al., 2025), which are mimicked here by transient CNO pulses (4 min). SST preferentially inhibits α-cells over β-cells, consistent with differential SSTR expression and signaling (Huising et al., 2025) providing a basis for asymmetric SST signaling within the islet. **(B)** Model-derived representative curves. SST was represented as a pulse-modulated signal (amplified by Gq or dampened by Gi). Insulin and glucagon outputs were modeled as the sum of SST-dependent inhibition and a delayed rebound/disinhibition component, with stronger SST sensitivity assigned to α-cells than β-cells (see Methods for equations). Rebound responses were triggered during SST decay phases following pulse offset. Model traces are illustrative and not fitted to individual experiments but constrained by experimentally measured amplitudes and delays (Figure 3) and are shown alongside experimental perifusion data to illustrate qualitative correspondence. **(C)** Model-derived net gains and hormone ratios. Central expected values are shown; values in parentheses indicate a fixed ±20% variation range used to illustrate robustness across plausible parameter choices (not statistical confidence intervals or fit uncertainty). **(D)** Experimental timeline in the anterior chamber of the eye. Mouse islets expressing SST-hM3Dq or SST-hM4Di were transplanted and animals randomly split into control and DREADD subgroups. **(E–F)** Plasma hormone levels in fasting and during feeding before and after chronic CNO (Days 3 & 15). **(G)** Net % change from untreated during feeding (Days 3 & 15 averaged; mean ± SEM). **(H)** Agreement between observed net gains during feeding and model-derived expectations (r² = 0.54, p = 0.0002). **(H1)** SST dynamically sets the INS/GCG balance: Gq promotes balance, Gi favors glucagon-skewed responses **(I)** Resolution of the human paradox: SST as dynamic modulator of islet plasticity. of high basal insulin coexisting with high somatostatin in type 2 diabetes (Fig. 7I).

### δ-cell post-inhibitory rebound transforms tonic inhibition into dynamic excitation, redefining intraislet plasticity

Data from isolated mouse islets (Figure 3) revealed that SST amplification (Gq) drives rebound excitation with balanced insulin/glucagon coupling favoring insulin, whereas SST dampening (Gi) produces disinhibition surges skewed toward glucagon (smaller INS/GCG ratio). These observations informed a data-informed phenomenological model of δ-cell pulse–rebound dynamics, where rebound responses were triggered during SST decay phases following pulse offset (**Figure 7A–C**). SST dynamics were represented as pulse-modulated signals (amplified by Gq or dampened by Gi, mimicking natural 5–15 min SST pulsatility), and insulin and glucagon outputs were modeled as the sum of SST-dependent inhibition plus a delayed rebound/disinhibition component, with stronger SST sensitivity assigned to α-cells than β-cells (see Methods). While our model is intended as a minimal phenomenological framework to capture experimentally observed dynamics rather than a fully mechanistic description of intracellular signaling, its formulation captures the key experimental features of δ-cell signaling, in which SST exerts both inhibitory and rebound-driving effects depending on its temporal dynamics. This accounts for the balanced rebound under Gq and the glucagon-skewed surge under Gi observed experimentally.

To test these expectations in a physiological context, mouse islets expressing SST-hM3Dq or SST-hM4Di were transplanted into the anterior chamber of the eye of immunodeficient mice and animals were randomly assigned to control or DREADD subgroups (**Fig. 7D**). Plasma hormones were measured in fasting and during feeding before and after chronic CNO administration in drinking water (Days 3 and 15; n=6 to 4 per group). One-tailed independent t-tests were used to compare each treated group with its own matched control cohort. In fasting, changes were modest (**Fig. 7E**). During feeding, a physiologically relevant state, Gq activation produced balanced increases in insulin and glucagon, whereas Gi inhibition drove a pronounced glucagon surge with relative sparing of insulin (**Fig. 7F, G**; p-values as indicated). This resulted in an insulin-favoring INS/GCG ratio under Gq and a glucagon-skewed ratio under Gi (**Fig. 7H1**).

Observed net gains during feeding showed significant agreement with model-derived central estimates (**Fig. 7H**; r² = 0.54, p = 0.0002). The model also recapitulated the positive INS–SST coupling observed under Gq amplification, providing a quantitative framework consistent with the elevated basal insulin and somatostatin levels and their positive correlation in human T2D islets (**Fig. 1E**; r² = 0.42, p = 0.0035). Plasma somatostatin could not be used for ratios because islet-derived SST contributes negligibly to the systemic pool.

Collectively, these results demonstrate that somatostatin is not merely a tonic inhibitor but a dynamic modulator that, through post-inhibitory rebound and differential sensitivity of α- and β-cells, confers plasticity and robustness to the endocrine pancreas, resolving the apparent paradox of high basal insulin coexisting with high somatostatin in type 2 diabetes (**Fig. 7I**).

These findings are summarized in the **Graphical Abstract**. Somatostatin, long considered a static inhibitor of secretion and growth, instead acts as a dynamic modulator that transforms inhibition into context-dependent excitation. In the islet, amplified SST pulses drive rebound excitation to promote balanced insulin and glucagon output during anabolic demand, while dampening produces glucagon-skewed surges for protection against hypoglycemia. Importantly, asymmetric δ-cell signaling, shaped by differential SSTR expression and downstream coupling, enables somatostatin to tune hormonal balance bidirectionally—favoring insulin- or glucagon-dominant states—establishing SST as a context-dependent regulator of physiological output rather than a static inhibitor. This pulse–rebound principle may extend to other somatostatin-regulated systems, including the brain, gut, and tumors, revealing a general principle in which SST signaling encodes diverse, context-dependent outcomes rather than enforcing uniform inhibition.

## DISCUSSION

This study was motivated by a striking and unexpected observation: islets from individuals with type 2 diabetes (T2D) exhibit concurrent elevation of both insulin and somatostatin secretion at baseline and across higher glucose concentrations, with tightly correlated profiles—contrasting with the decoupled dynamics typically observed in healthy islets. In non-diabetic islets, basal hormone output is low, and glucose stimulation produces an asymmetric response: insulin rises steeply while somatostatin increases modestly, yielding a high insulin:somatostatin ratio that reflects β-cell dominance and restrained δ-cell tone (Kailey et al., 2012; Briant et al., 2018; Vergari et al., 2019; 2020). This asymmetry is essential for paracrine regulation of α- and β-cells and for systemic processes such as hepatic insulin clearance (Ipp et al., 1987; Strowski & Blake, 2008). By contrast, the T2D phenotype—a lower insulin:somatostatin ratio with preserved or elevated δ-cell output (Rorsman & Huising, 2018; Vergari et al., 2020; Oduori et al., 2020; Yang et al., 2025)—presents a paradox: how can δ-cell “dominance” coexist with hyperinsulinemia and inappropriate glucagon secretion in the setting of impaired β-cell competence?

Our findings offers a mechanistic explanation to this paradox by demonstrating that δ-cell activity can paradoxically prime and synchronize excitatory responses through post-inhibitory rebound (PIR), also referred to as post-inhibitory excitation (PIE). This framework provides a unifying explanation for previously discordant observations in δ-cell biology. Chemogenetic and physiological manipulations showed that transient δ-cell activation suppresses insulin and glucagon but is consistently followed by rebound peaks. Such biphasic dynamics—acute inhibition followed by overshoot excitation—are hallmarks of PIR observed across excitable systems including thalamocortical circuits, brainstem respiratory networks, and cardiac pacemaker cells (Smith et al., 1991; Steriade et al., 1993; DiFrancesco, 1993; McCormick & Bal, 1997). PIR thus provides a mechanistic framework connecting our human islet data, mouse DREADD ex vivo perifusion dynamics and in vivo experiments, together with earlier studies showing rebound hormone release after somatostatin withdrawal (Leblanc et al., 1975; Ward et al., 1975; Bratusch-Marrain et al., 1981). While rebound responses have been described in excitable systems, their role as a primary organizing principle of endocrine coordination has not been previously recognized.

Several convergent mechanisms underlie PIR in α- and β-cells, largely through Gi/o receptor signaling, the canonical pathway downstream of SSTRs. First, Gi/o activation suppresses cAMP production and downstream effectors such as PKA and EPAC, reducing granule priming, exocytotic competence, and ion channel facilitation (Kagimoto et al., 1994; Doyle & Egan, 2007; Roed et al., 2012; Tengholm & Gylfe, 2017; Yu et al., 2019; Dyachok et al., 2021; Dickerson et al., 2022). Relief of inhibition allows rapid cAMP recovery with transient overshoot, amplifying Ca²⁺-triggered secretion. Second, Gi/o signaling induces hyperpolarization not only via GIRK channels but also through Src-family kinase–dependent Na⁺/K⁺-ATPase (NKA) activation, which suppresses excitability; rebound occurs when reduced NKA activity permits post-inhibitory depolarization and synchronized Ca²⁺ entry (Dickerson et al., 2022). Third, Ca²⁺ handling contributes: inhibition favors ER refilling, and upon release, store-operated Ca²⁺ entry (SOCE) and Ca²⁺-induced Ca²⁺ release generate amplified cytosolic transients (Berts et al., 1997; Chen et al., 2003; Tian et al., 2011; Rorsman et al., 2012; Sabourin et al., 2015; Klec et al., 2019; Zhang et al., 2020). Finally, δ-cell somatostatin release is pulsatile and phase-locked to islet Ca²⁺ oscillations; release and withdrawal therefore synchronize rebound responses across α- and β-cells (Hauge-Evans et al., 2009; Zhang et al., 2020; Ren et al., 2022).

The rebound hypothesis also offers a mechanistic explanation for sustained insulin and somatostatin secretion in T2D. Chronic δ-cell activation—driven by hyperglycemia or altered paracrine feedback—may foster δ-cell hyperexcitability or enhanced tonic tone, sustaining insulin secretion through repetitive rebound cycles. PIR-like behavior in β-cells has been proposed to arise from “hyperexcitation” during prolonged glucose exposure, a state thought to underlie reversible insulin secretory defects (Nichols & Remedi, 2012). This may explain early hyperinsulinemia preceding β-cell failure (Weir & Bonner-Weir, 2004) and reconcile prior discrepancies: somatostatin may acutely suppress secretion while promoting pathological hypersecretion over longer timescales (Omar-Hmeadi et al., 2020; Yang et al., 2025).

Rebound dynamics extend to glucagon. Somatostatin inhibits α-cells as part of normal paracrine control (Gromada et al., 2007; Vergari et al., 2020). Yet relief of inhibition can generate amplified glucagon bursts, consistent with α-cell expression of Gi/o-coupled SSTR2 and α₂-adrenergic receptors (Salehi et al., 2007; Hellman et al., 2009; Zhang et al., 2013). Hyperpolarization via NKA reduces Ca²⁺ influx, and rebound depolarization reopens Ca²⁺ channels, triggering synchronized glucagon release. Pulsatile somatostatin may thus paradoxically potentiate glucagon secretion, explaining inappropriate dual hypersecretion in T2D.

Frequent δ-cell activation followed by inhibition amplifies rebound probability and magnitude, producing net rises in insulin and glucagon measured in vivo, including in high-fat-diet models. δ-cell “dominance” therefore reflects active shaping of secretion dynamics, not static suppression. DREADD-based manipulations support this model: selective δ-cell activation suppressed insulin and glucagon followed by rebound peaks resembling human studies (Leblanc et al., 1975; Ward et al., 1975; Bratusch-Marrain et al., 1981). Conversely, δ-cell inhibition triggered glucagon disinhibition and smaller insulin rebounds, consistent with distinct α- and β-cell sensitivities and receptor expression (Kailey et al., 2012; Gromada et al., 2007). In vivo, δ-cell activation enhanced glucose tolerance in transplantation models; although systemic insulin and glucagon levels rose—particularly relative to somatostatin—the insulin:glucagon ratio remained balanced toward insulin dominance. By contrast, δ-cell inhibition in vivo markedly increased circulating glucagon and shifted the ratio toward glucagon dominance.

Re-analysis of raw β-cell Ca²⁺ recordings from a prior collaborative optogenetic study (Arrojo e Drigo et al., Nat Commun 10, 3700, 2019), in which δ-cells were photoactivated in the anterior chamber of the eye model, now reveals that the initial short transient inhibition of β-cell Ca²⁺ activity is followed by a net PIR-like increase, with further enhancement upon a second light pulse. This previously unappreciated net excitatory effect explains the unexpected systemic hypoglycemia and elevated plasma insulin we observed in those early experiments and aligns with the PIR mechanism demonstrated here across chemogenetic, ligand, and in vitro approaches.

These results reveal that somatostatin-driven Gi/o signaling generates structured sequences of inhibition followed by post-inhibitory rebound, reconciling the paradoxical co-elevation of insulin and somatostatin in T2D. The insights extend beyond islets: PIE appears as a conserved regulatory motif across neuroendocrine and visceral systems, with implications for therapeutic targeting. Future directions include dissecting NKA and SOCE contributions to PIR, defining δ-cell timing changes under chronic metabolic stress, and testing temporally patterned δ-cell modulation as a strategy to restore physiological hormone rhythms and improve glycemic control.

Several additional considerations refine the PIR framework. Receptor subtype and coupling are central: in human α- and β-cells, SSTR5 and SSTR2 predominate and signal through Gi/o (Strowski and Blake, 2008; Vergari et al., 2019), dictating whether somatostatin action leads to GIRK-mediated hyperpolarization, NKA activation, or cAMP suppression. Importantly, recent independent work has defined receptor-specific SST signaling properties that further support asymmetric control of α- and β-cells (Huising and colleagues, 2025). Variation in expression across species or disease states likely underlies differences in rebound amplitude and timing (Kailey et al., 2012; Roed et al., 2012). Other paracrine and hormonal cues also shape PIR; in our study, both urocortin-3 and ghrelin enhanced insulin rebound by modulating δ-cell activity and somatostatin release (van der Meulen et al., 2015; DiGruccio et al., 2016). Rebound dynamics may be further tuned by sex and metabolic state, as seen in glucagon sensitivity. Beyond these, chronic disease remodeling—altered receptor expression, ion channel composition, gap junction connectivity (Briant et al., 2018), and ER/SOCE machinery—reshapes inhibitory tone and rebound capacity, contributing to the heterogeneity of T2D.

These findings emphasize a dynamic consequence of δ-cell modulation beyond static inhibitory models, suggesting that timed δ-cell engagement may be harnessed to improve postprandial glucose control. Systemically, δ-cell activity influences hepatic insulin clearance and hormone kinetics (Blackard & Nelson, 1970; Efendic et al., 1975; 1976; Sherwin et al., 1976; Bonora et al., 1983; Ipp et al., 1987). Thus, improved glucose tolerance following δ-cell activation likely reflects both intra-islet PIR-mediated insulin augmentation and reduced hepatic clearance. Similarly, PIR-mediated glucagon enhancement may affect hepatic glucose production, exacerbating hyperglucagonemia and hyperglycemia in T2D. These systemic effects underscore the translational importance of δ-cell signaling, suggesting therapies targeting somatostatin pathways could modulate islet secretion, SSTR re-sensitization, and hormonal clearance dynamics.

Limitations include incomplete understanding of NKA regulation by GPCR–SFK pathways in α-versus β-cells, a focus on acute dynamics with minimal longitudinal data, species differences in islet architecture and receptor expression, and the broader δ-cell secretome and intercellular interactions that may influence network timing and PIR. Despite these caveats, human perfused preparations and infusion studies suggest conserved rebound phenomena amenable to translational exploration (Leblanc et al., 1975; Ward et al., 1975; Bratusch-Marrain et al., 1981).

In conclusion, δ-cells function as dynamic temporal orchestrators of islet excitability, shaping insulin and glucagon rhythms through rebound excitation. In healthy islets, transient δ-cell activation and somatostatin release synchronizes α- and β-cell activity for appropriate pulsatility (Salehi et al., 2007; Briant et al., 2017). In T2D, chronic δ-cell activation or SSTR desensitization may drive maladaptive rebounds, contributing to dysregulated glycemia.

Analogous to inhibitory interneurons in neural networks, δ-cells suppress and coordinate excitatory output (Isaacson & Scanziani, 2011). Beyond islets, somatostatin-driven PIE represents a conserved regulatory principle across neuroendocrine, gastrointestinal, and sensory systems (Barbieri et al., 2013; Corleto et al., 2006; Hagströmer et al., 2006; Huang et al., 2018). Understanding this temporal logic opens avenues for δ-cell modulation in diabetes and broader PIE-based interventions in cancer, inflammation, and sensory physiology.

These findings redefine the role of somatostatin and δ-cells in the endocrine pancreas. Rather than acting as a static brake, SST operates as a dynamic modulator that transforms inhibition into excitation through post-inhibitory rebound. When δ-cell activity is amplified, rebound responses generate balanced, robust secretion of both insulin and glucagon, enabling the islet to meet high physiological demands while preserving glycemic stability. Conversely, when δ-cell tone is reduced, disinhibition produces glucagon-skewed output, providing a protective mechanism against hypoglycemia. This mechanism explains the paradoxical coexistence of elevated basal insulin and somatostatin in human type 2 diabetes and reveals how δ-cells confer plasticity and robustness to the islet micro-organ.

Although δ-cells serve as a physiological hub for this mechanism, similar rebound dynamics can be elicited through other intra-islet inhibitory interactions. However, the δ-cell is uniquely positioned for this role: it is the most densely innervated endocrine cell type in both mouse and human islets (Rodriguez-Diaz et al., 2011), making it an ideal integrator of central, local paracrine, and vascular signals. Many signals previously viewed as purely inhibitory may instead act through δ-cells to fine-tune somatostatin pulses, thereby enhancing rather than diminishing overall islet output when needed. The δ-cell may thus function as a tunable “atrium” of the islet micro-organ, allowing the pancreas to orchestrate a precise hormonal symphony in response to systemic needs (see Graphical Abstract).

Importantly, asymmetric SST signaling—arising from differential SSTR expression and downstream coupling in α- and β-cells—enables δ-cells to tune hormonal balance bidirectionally. Stronger inhibition of α-cells relative to β-cells favors balanced rebound under conditions of increased SST, whereas reduced SST tone permits glucagon-dominant disinhibition. This asymmetry provides a mechanistic basis for the flexible control of hormone ratios across physiological states and is consistent with recent independent reports defining receptor-specific SST signaling properties (Huising and colleagues, 2025). Together, these findings support a revised model of intra-islet paracrine crosstalk in which δ-cells do not simply impose inhibition but dynamically encode future excitation through pulse-dependent rebound mechanisms.

Together, these discoveries establish a revised model of intra-islet paracrine crosstalk in which δ-cell–driven temporal dynamics, rather than tonic inhibition, determine endocrine coordination. In this framework, inhibition is not merely suppressive but instructive, encoding future hormone output through rebound dynamics. While rebound responses have been described in excitable systems, their role as a primary organizing principle of endocrine signaling has not been previously recognized. This framework provides a unifying explanation for prior observations in δ-cell biology that were difficult to reconcile under static inhibitory models.

More broadly, we believe the main contribution of our work here is the suggestion that somatostatin signaling may encode context-dependent outcomes through its temporal dynamics rather than acting solely as a tonic inhibitory signal. While this study establishes this principle in the pancreatic islet, similar paradoxes in systems where SST is present—including neural circuits, gastrointestinal regulation, and proliferative or inflammatory contexts—raise the possibility that dynamic SST signaling represents a more general mechanism for modulating tissue excitability and homeostasis. In this view, SST should not be considered merely suppressive, but rather as a regulator capable of tuning system output across physiological states.

This dynamic framework has important implications for disease. Context-dependent SST signaling may contribute to inflammation, tumor progression, and metabolic dysfunction in ways that are not captured by static inhibitory models. Notably, recent reports of δ-cell remodeling and altered β–δ coupling in human T1D (Hill et al., 2024; Tegehall et al., 2025), together with refined understanding of SSTR subtype specificity (Huising and colleagues, 2025), align with the hypersecretory δ-cell phenotype we and others have observed. Our identification of PIR provides a mechanistic framework that helps reconcile these observations and offers a unifying explanation for the seemingly contradictory roles of somatostatin across physiological and pathological states.

By establishing δ-cells and somatostatin as central regulators of endocrine plasticity, this work opens new avenues for understanding and therapeutically targeting hormone balance in diabetes and beyond. More broadly, defining how SST-dependent rebound signaling operates across tissues may reveal a general principle of biological regulation that has remained largely underappreciated.

## Supporting information

Supplementary FIGURES with legends and Suppelmentary Tables

## STAR Methods

### RESOURCE AVAILABILITY

#### Lead Contact

Further information and requests for resources and reagents should be directed to and will be fulfilled by the Lead Contact, **Dr. Rayner Rodriguez-Díaz**.

#### Materials Availability

All unique mouse lines and reagents generated in this study are available from the Lead Contact upon reasonable request and execution of a material transfer agreement (MTA).

#### Data and Code Availability

All data supporting the findings of this study are provided in the paper and Supplementary Information. Raw perifusion time-course data and in-vivo metabolic datasets are available from the Lead Contact upon reasonable request. No custom code was used.

## EXPERIMENTAL MODEL AND SUBJECT DETAILS

### Human Islet Donors

Human pancreatic islets were obtained from non-diabetic (ND) and type 2 diabetic (T2D) donors via the Integrated Islet Distribution Program and the Diabetes Research Institute Human Cell Processing Core (University of Miami) or purchased from Prodo Laboratories. Donor metadata (age, sex, BMI, HbA1c) are listed in Table S1. Islets were cultured overnight in CMRL-1066 (5.5 mM glucose) supplemented with 10% FBS, 2 mM L-glutamine, and 100 U/mL penicillin/streptomycin before experiments. Preparations with >80% viability and <20% exocrine contamination were used.

#### Human research compliance

De-identified human tissue was used in accordance with institutional guidelines and IRB policies for non-human subjects research/waived consent.

### Mouse Models

For diet-induced obesity studies, C57BL/6J mice were maintained on standard chow (RD) or 45% high-fat diet (HFD; Research Diets D12451) for 30 days prior to islet isolation.

For chemogenetics, Cre drivers were crossed to Cre-dependent DREADD reporter lines to target specific endocrine subsets:

- **β-cell activation:** Ins2-Cre (JAX 003573) × ROSA-hM3Dq-HA-citrine (JAX 026220) → INS-DREADD-Gq
- α-cell activation: Gcg-iCre (JAX 030663) × ROSA-hM3Dq-HA-citrine (JAX 026220) → GLUCAGON-DREADD-Gq
- δ-cell activation: Sst-Cre (JAX 013044) × ROSA-hM3Dq-HA-citrine (JAX 026220) → SOMATOSTATIN-DREADD-Gq
- δ-cell inhibition: Sst-Cre × ROSA-hM4Di-HA-citrine (JAX 026219) → SOMATOSTATIN-DREADD-Gi

Additional δ-cell Gq lines used for imaging/reporters:

- **SOMATOSTATIN-DREADD-Gq; GCaMP3:** (Sst-Cre × ROSA-hM3Dq) × ROSA-GCaMP3 (JAX 029043)
- **SOMATOSTATIN-DREADD-Gq-mCherry:** Sst-Cre × ROSA26-hM3Dq-mCherry (JAX 026943)

Both sexes (2–3 months) were used unless stated otherwise. Genotypes were confirmed by PCR.

#### Animal ethics

Procedures were approved by the University of Miami IACUC and complied with NIH guidelines.

#### Sex as a biological variable

Sex was recorded for all donors and mice; analyses stratified by sex are reported where relevant.

## METHOD DETAILS

### Mouse Islet Isolation and Culture

Islets were isolated by collagenase P digestion, purified by density gradient, and hand-picked. For diet studies, islets were cultured 4 h before perifusion. For chemogenetic studies, islets were cultured overnight in RPMI-1640 (11 mM glucose, 10% FBS, antibiotics) prior to perifusion or transplantation.

### Perifusion Assays and Chemogenetic Manipulations (in vitro)

Dynamic secretion was measured in parallel chambers (Biorep) at 37 °C with constant flow **0.06 mL/min** of KRB (in mM: 115 NaCl, 5 KCl, 24 NaHCO₃, 2.5 CaCl₂, 1 MgCl₂, 10 HEPES; 0.1% BSA; pH 7.4). Islets (**150 IEQ**/chamber) equilibrated 30 min, then underwent glucose ramps:

- **Mouse islets:** 3–16 mM or 3→16→1 mM
- **Human islets:** 3→11 mM or 3→11→1 mM

### Chemogenetic stimulation

For cell-specific activation, DREADD-Gq islets received sequential **4-min** CNO pulses (1, 10, 100 µM for mouse, 2 & 10 µM for human) spaced **8 min** apart at either 16 mM or 1 mM glucose. For δ-cell inhibition, SOMATOSTATIN-DREADD-Gi islets received identical CNO pulse trains. Fractions were collected every **2 min**.

Insulin and glucagon were quantified by ELISA (Mercodia); somatostatin by ELISA (Phoenix Pharmaceuticals). PIR insulin was assessed at **3, 7, and 16 mM** glucose and two sequential 4-min 2.5 µM CNO pulses spaced **8 min** apart. Where indicated, **Cyclosomatostatin (500 nM)** was included at 16 mM glucose to test SSTR dependence of **post-inhibitory rebound (PIR)** insulin responses. Experiments concluded with KCl (depolarization, 25 mM) to assess maximal secretory capacity.

### Expected pharmacology (summary)

DREADD-Gq produced dose-dependent homeotypic secretion (insulin, glucagon, or somatostatin) at permissive glucose, with occasional desensitization at 100 µM CNO; DREADD-Gi reduced somatostatin by ∼60% and disinhibited glucagon.

### Ligand Stimulation Studies (Endogenous δ-cell Activation)

Perifusion assays at **16 mM** glucose tested **Urocortin-3 (Ucn3)** and **Ghrelin (Ghr)** at **2.5 nM** and **25 nM** (pulses for mouse islets) or 1**2.5 nM** and **100 nM** (pulses for human islets) Both ligands increased somatostatin and reproduced the biphasic “inhibition-then-rebound” insulin secretion profile.

### Immunocytochemistry and Confocal Imaging

Pancreata and isolated islet adsorbed to glass slides were fixed in 4% paraformaldehyde. Tissue sections were cryoprotected, and cryosectioned (40 µm). Sections were stained with anti-HA (Cell Signaling 3724), anti-insulin (Abcam ab7842), anti-glucagon (Sigma G2654), anti-somatostatin (Abcam ab53165), Alexa Fluor secondaries, and DAPI. Confocal imaging (Leica SP8) confirmed cell-type specificity and membrane localization of HA-tagged DREADDs.

### Adenoviral transduction of human δ-cells

Human islets from non-diabetic donors were transduced with a replication-deficient Ad5/F35 chimeric adenoviral vector expressing hM3D(Gq) fused to mCherry under the control of the human somatostatin (huSST) promoter (AdV-Sst-DREADD-Gq, VectorBuilder ID VB221116-1460qpe) at a multiplicity of infection (MOI) of 50. The Ad5/F35 backbone was selected to enhance transduction efficiency in primary human islets, which exhibit low coxsackievirus and adenovirus receptor (CAR) expression. Full vector design, map, and sequence are provided in Supplementary Information 1. Islets were incubated for 60–160 h post-transduction in CMRL-1066 medium supplemented with 10% FBS. δ-cell specificity and transduction efficiency were confirmed by immunocytochemistry (Supplementary Figure S9). Non-transduced islets from the same preparations served as controls.

### Calcium imaging

Transduced islets were loaded with the fluorescent Ca²⁺ indicator Calbryte 520 for 1 h at 37 °C and imaged using confocal microscopy. Chemogenetic activation was induced by application of clozapine-N-oxide (CNO). mCherry-positive regions (δ-cells) were analyzed for intracellular Ca²⁺ dynamics.

### Perifusion assays

(chemogenetic activation in human islets). For experiments involving chemogenetic activation in transduced human islets, CNO was applied as 4-min pulses (2 µM and 10 µM) during defined glucose conditions (3 mM, 16 mM, or 1 mM). Experiments concluded with KCl depolarization to assess maximal secretory capacity.

### In Vivo Islet Transplantation and Engraftment

To isolate δ-cell contributions to systemic glucose control, **200 IEQ** from F1 donors were transplanted into the anterior chamber of the eye of **female athymic nude mice (10 weeks)** rendered diabetic with **streptozotocin (200 mg/kg, i.v.)**. Normoglycemia returned within 30–40 days. Recipients were grouped by donor genotype (SOMATOSTATIN-DREADD-Gq, SOMATOSTATIN-DREADD-Gi, or Sst-Cre controls) and by donor sex. Mice were maintained ≥90 days before testing. **In vivo δ-cell Ca²⁺ imaging.** A subset received contralateral grafts from **SOMATOSTATIN-DREADD-Gq; GCaMP3** mice. Fluorescence responses to sequential i.p. CNO (**0.5 and 5 mg/kg**) were recorded with a custom laser-scanning setup and analyzed in ImageJ.

### In vivo optogenetic activation of δ-cells and β-cell Ca²⁺ imaging (Figure 5)

SST-ChR2 islets (or δ-cell-ablated controls) were transplanted into the anterior chamber of the eye of STZ-diabetic nude mice. After vascularization and engraftment (confirmed by restoration of normoglycemia), blue-light stimulation (4×15 s + 4×60 s pulses) was delivered to activate δ-cells while monitoring systemic glucose and plasma insulin. For **β-cell Ca²⁺ imaging**, a subset of grafts expressed GCaMP6s in β-cells. Data shown in Figure 5I–J were re-analyzed from raw recordings originally acquired in collaboration with R. Arrojo e Drigo (Arrojo e Drigo et al., Nat Commun 10, 3700, 2019) and demonstrate net increase in β-cell Ca²⁺ activity after δ-cell photoactivation, with further enhancement upon a second light pulse.

### Metabolic Testing in Transplanted Mice. Testing was performed under: postprandial

(2 h after feeding onset; ZT14), **short fasting** (2 h), or **long fasting** (16 h overnight). Glycemia (OneTouch Ultra) and plasma hormones (ELISA) were measured.

### CNO Administration

For **acute** δ-cell manipulation, **CNO 5 mg/kg i.p.** was given **30 min** before tests; controls received saline. For **short-chronic** manipulation, CNO was provided in drinking water for 7 days (estimated **5 mg/kg/day**, assuming ∼4 mL/mouse/day).

### Intraperitoneal Glucose Tolerance Test (IPGTT)

Following 16 h fasting, mice received **2 g/kg** glucose i.p. Blood glucose was measured at -30, 0, 15, 30, 45, 60, 90, and 120 min.

Hormone levels were estimated at -30, 0, 15, and 120 min. Each animal underwent two sessions with crossover (CNO vs saline).

### Insulin Tolerance Test (ITT)

After 2 h fasting, mice received **0.75 IU/kg** insulin i.p. following saline or CNO pretreatment. Blood glucose measures mirrored the IPGTT, while glucagon was estimated at -30, 0 and 30 min.

### Observed pharmacodynamics (summary)

Acute δ-cell activation produced a transient hypoglycemia accompanied by elevated plasma insulin. During ITT, both activation and inhibition of δ-cells resulted in reduced plasma glucagon levels compared with controls. Notably, despite this suppression, the overall glucose counterregulatory response to insulin-induced hypoglycemia remained unaffected.

## QUANTIFICATION AND STATISTICAL ANALYSIS

### Perifusion Traces

Perifusion profiles are presented preferentially as absolute hormone concentrations in effluent fractions collected every 2 min (temporal resolution: 2 min). When indicated in figure panels, traces were normalized to the maximal KCl response or baseline-corrected.

For comparisons within the same islet isolation, minor adjustments were applied at loading to ensure comparable total protein content across chambers.

Peak amplitude and area under the curve (AUC) were calculated for each stimulus. Post-inhibitory rebound (PIR) insulin secretion was defined as the AUC above baseline following CNO withdrawal, or relative to no-DREADD control islets subjected to identical stimulation protocols.

### Phenomenological modeling of δ-cell pulse–rebound dynamics

A data-informed phenomenological model was developed to summarize the acute hormone dynamics elicited by δ-cell hM3Dq (Gq) or hM4Di (Gi) manipulation and to predict chronic in vivo outcomes. SST output was represented as a pulse-modulated signal that mimics natural 5–15 min pulsatility: SST(t) = S₀ + s·A·u(t) where S₀ is basal SST tone, rebound responses were triggered during SST decay phases following pulse offset (τ ≈ 2 min), A is the magnitude of chemogenetic modulation, u(t) is a binary pulse function representing the timing of CNO stimulation, and s = +1 for Gq and s = −1 for Gi.

Insulin and glucagon responses were modeled as: INS(t) = I₀ − w_I·(SST(t)−S₀) + R_I(t), GCG(t) = G₀ − w_G·(SST(t)−S₀) + R_G(t) where w_I and w_G are inhibitory weights (w_G > w_I to encode stronger SST sensitivity of α-cells) and R_I(t) and R_G(t) are delayed rebound/disinhibition terms triggered by relief of SST-dependent inhibition. Delayed responses were modeled as exponentially decaying kernels initiated at SST pulse offset: R_i(t) = r_i · H(t − t_off) · exp(−(t − t_off)/τ_i) where H is the Heaviside step function, r_i is rebound gain, and τ_i is the decay constant (τ ≈ 2 min from experimental delays in Figure 3).

Model parameters were chosen to reproduce the main temporal and amplitude features observed in Figure 3 (e.g., SST peak amplitudes 120–220%, rebound delays 3–6 min, net gains). Thus, model traces are illustrative and not fitted to individual experiments but constrained by experimentally measured amplitudes and delays (Figure 3). Central model-derived gains are reported together with a fixed ±20% variation range for illustrative robustness (not derived from parameter fitting or statistical uncertainty but used to illustrate robustness across plausible parameter ranges). Agreement with in vivo feeding responses was assessed by linear regression.

Importantly, this model is intended as a mechanistic abstraction capturing the qualitative and semi-quantitative features of δ-cell-driven hormone dynamics rather than a fully parameter-fitted predictive model.

**Statistical analyses** were performed in **GraphPad Prism 10**. Two-group comparisons used paired or unpaired two-tailed **t-tests** as appropriate. Multi-factor experiments used **two-way ANOVA** with **Šidák** post-hoc correction. Data are mean ± SEM; **p < 0.05** was considered significant.

### Randomization and blinding

For in-vivo tests, animals were randomly assigned to treatment order (CNO vs saline crossover). Sample processing and ELISA quantification were performed with investigators blinded to group when feasible.

### Exclusion criteria

Pre-specified criteria included perifusion duplicate failure or abnormally high or low hormone concentrations related to the data. Also, surgical failure to engraft (no reversion of hyperglycemia by day 40) or death during the experimental protocol. excluded cases are reported in figure legends where applicable.

## KEY RESOURCES TABLE

(To be completed at final submission; will list antibodies, mouse strains/lines, ligands, CNO source, ELISAs with catalog numbers, software.)

## Article Information

### Acknowledgments

This work was supported by NIH grants R01DK124527 & R21DK114418 (to R.R.-D.) and 6048/2013SW, Raising Foundation Diabetes Wellness Network Sweden (to R.R-D). We thank the teams led by Alejandro Caicedo, Joana Almaça, Manuel Blandino, and Ernesto Bernal for their invaluable logistical and scientific support. We are deeply grateful to Per-Olof Berggren and Barbara Corkey for their mentorship, critical review of the manuscript, and continued encouragement and support.

### Duality of Interest

The authors declare no competing interests.

### Author Contributions

A.T-G, D.H-R, O.A, E.P, S.C-B, L.M-C, M.S, R.AeD & R.R-D performed the experiments. A.T-G., D.H-R, P.O.B, R.AeD and R.R-D analyzed the results. R.R-D. conceived the study. A.T-G., D.H-R, O.A, R.AeD and R.R-D. designed the experiments and interpreted the results. R.R-D. wrote the paper. All authors read and approved the final version of the paper. A.T.G, D.H.R and R.R-D. are guarantors of the work and take responsibility for the integrity and accuracy of the data and data analysis. Further information and requests for resources, reagents, and data should be directed to and will be fulfilled by A.T-G, D.H-R and R.R-D.

## Summary of Experimental Logic

| Experimental Arm | Aim | Manipulation | Readouts |
| --- | --- | --- | --- |
| Human islet perfusion | Assess parallel insulin–somatostatin dynamics | Glucose steps | Hormone output & secretion kinetics |
| Mouse RD vs HFD islet perfusion | Test basal/stimulated hypersecretion | Diet manipulation | Hormone output |
| $\alpha/\beta/\delta$ DREADD-Gq / $\delta$ DREADD-Gi characterization / intra islet cell communication | Dissect α–β–δ cell crosstalk in the islet & dissect δ-cell effects on $\alpha/\beta$ cells | CNO pulse trains / chemogenetics | PIR and rebound dynamics, co-stimulation and disinhibition surges. New Model for Paracrinicity |
| Glucose, Cyclosomatostatin Ligand (Ucn3, Ghr) | Test endogenous δ-cell-driven PIR / Physiological context | Pulsed native ligands / antagonist | Rebound insulin secretion |
| Human AdV-Sst-DREADD-Gq transduction | δ-cell specific chemogenetic activation in human islets | AdV-Sst-DREADD-Gq transduction + CNO/perifusion | Immunocytochemistry, triple-hormone secretion. Rebound dynamics. |
| Human AdV-Sst-DREADD-Gq transduction | Cellular mechanism | AdV-Sst-DREADD-Gq + CNO, Calbryte520 | Intracellular $\text{Ca}^{2+}$ transients (bulk imaging), Rebound insulin secretion |
| Intraocular graft model chemogenetic manipulation | Evaluate systemic relevance of specific δ-cell manipulation | CNO ip injected δ-cell Gq/Gi (acute) | Glycemia, plasma hormones, IPGTT/ITT |
| Intraocular graft model chemogenetic manipulation | Hormone balance | CNO in the water δ-cell Gq/Gi (chronic) | Hormone balance |
| In vivo optogenetic δ-cell activation (α-cell control) | Orthogonal validation of δ-cell-driven PIR in vivo | SST-ChR2 & GCG-ChR2 photoactivation in ACE grafts (blue light pulses) | Systemic glucose, plasma insulin |
| In vivo optogenetic δ-cell activation (α-cell control) | Targeted $\delta/\alpha$-cell ablation | Diphtheria-Toxin-Targeted $\delta/\alpha$-cell ablation | Readout specificity |
| In vivo optogenetic δ-cell activation + β-cell $\text{Ca}^{2+}$ imaging | In vivo δ-cell light activation | In vivo δ-cell activation (blue light) & β-cell-GCamp6 | Intracellular $\text{Ca}^{2+}$ transients (β-cell imaging) |
| Phenomenological modeling of δ-cell pulse–rebound dynamics | Synthesize acute data and predict chronic outcomes | Data-informed pulse-modulated SST model | Net hormone gains, INS/GCG ratio shifts |

## References (Cell style, numbered for appearance in text)

### Declaration of generative AI and AI-assisted technologies in the manuscript preparation process

During the preparation of this work the author(s) used ChatGPT & Grok to improve the writing process to improve the clarity and readability and language of the manuscript. After using this tool/service, the author(s) reviewed and edited the content as needed and take(s) full responsibility for the content of the published article.

## Notes

### Competing Interest Statement

The authors have declared no competing interest.

### Summary of Updates

-Main figures updated -Supplemental figure updated -Results section updated to match the figures -Discussion language modified to interpret the results

