## Supplementary FIGURES with legends and Suppelmentary Tables for "Post-Inhibitory Rebound by δ-Cells Converts Inhibition into Excitation and Confers Islet Plasticity"

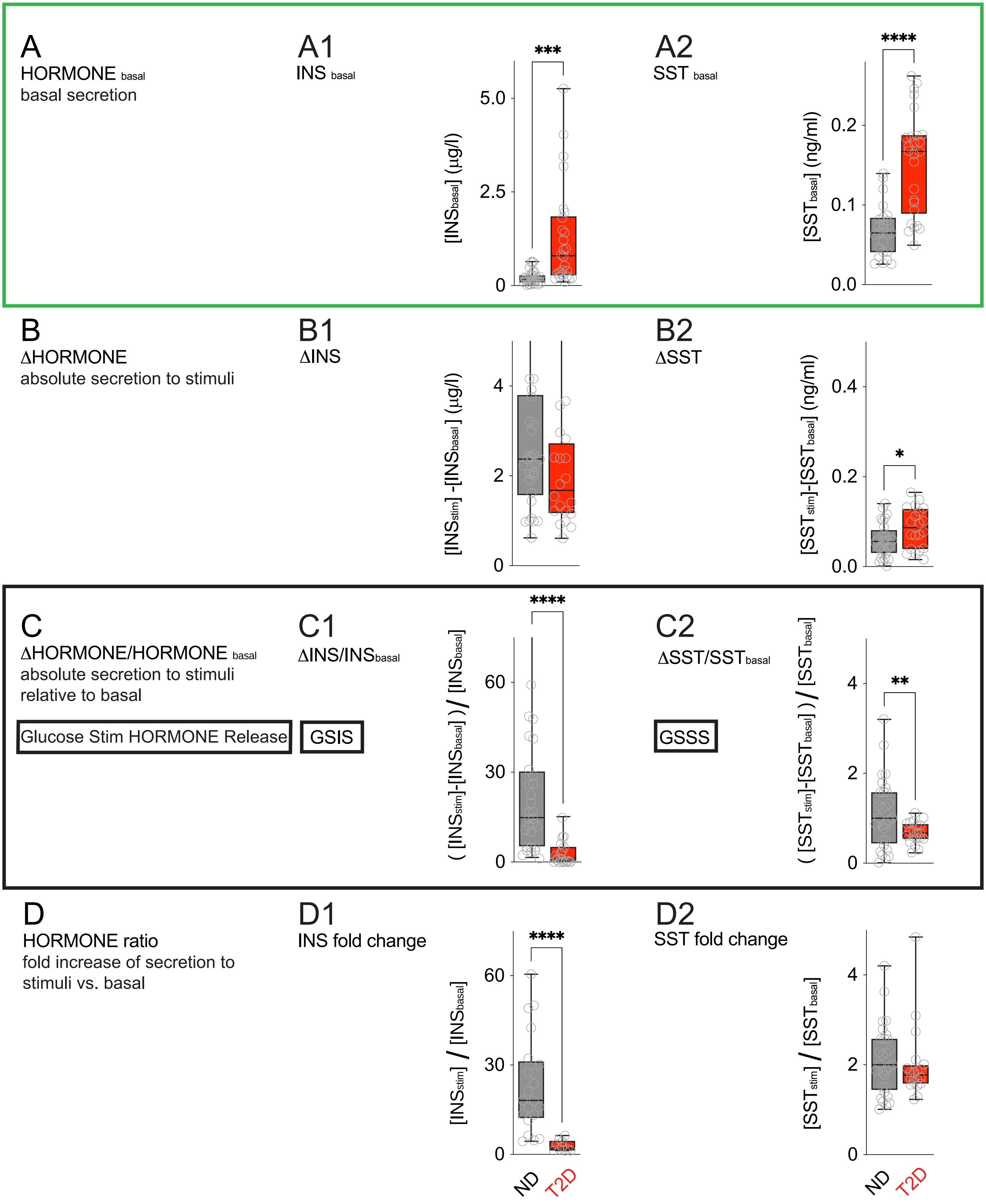

A
HORMONE basal
basal secretion
A1
INS basal
[INSbasal] (µg/l)
***
A2
SST basal
[SSTbasal] (ng/ml)
****
B
ΔHORMONE
absolute secretion to stimuli
B1
ΔINS
[INSstim] -[INSbasal] (µg/l)
B2
ΔSST
[SST stim]-[SSTbasal] (ng/ml)
*
C
ΔHORMONE/HORMONE basal
absolute secretion to stimuli
relative to basal
Glucose Stim HORMONE Release
GSIS
C1
ΔINS/INSbasal
( [INSstim]-[INSbasal] ) / [INSbasal]
****
C2
ΔSST/SSTbasal
GSSS
( [SST stim]-[SSTbasal] ) / [SSTbasal]
**
D
HORMONE ratio
fold increase of secretion to
stimuli vs. basal
D1
INS fold change
[INSstim] / [INSbasal]
****
D2
SST fold change
[SST stim] / [SSTbasal]
ND
T2D

**Figure S1. Type 2 Diabetic Human Islets Exhibit Elevated Basal Hormone Secretion and Impaired Glucose-Stimulated Hormonal Responses** (related to Figure 1). Islets of Langerhans isolated from individuals with type 2 diabetes (T2D) exhibit elevated basal hormone secretion, resulting in a diminished glucose-stimulated hormonal response (11 mM glucose) compared to islets from non-diabetic (ND) donors. This dysregulation is evident in both insulin and somatostatin release dynamics.

**(A, A1, A2)** Basal secretion levels of insulin and somatostatin, respectively, measured during perifusion with 3 mM glucose (designated as $HORMONE_{basal}$, basal background). Islet preparations were obtained from ND (n = 15; HbA1c < 5.6; BMI < 27) and T2D (n = 15; HbA1c > 6.0) human donors.

Glucose-stimulated secretion in response to 11 mM glucose was quantified using three analytical approaches:

**(B, B1, B2)** *Absolute change from basal* ($\Delta HORMONE = max[HORMONE_{stim}] - [HORMONE_{basal}]$: Quantifies the net increase in hormone secretion from baseline, indicating the magnitude of glucose-induced secretion.
**(C, C1, C2)** *Relative change from basal* ($\Delta HORMONE/HORMONE_{basal} = ( [HORMONE_{stim}] - [HORMONE_{basal}]) / [HORMONE_{basal}]$. Represents the glucose-stimulated insulin secretion (GSIS) and glucose-stimulated somatostatin secretion (GSSS) indices by normalizing the absolute change to the basal level. It is highlighted in the Figure because this is the generalized way investigators report similar data.
**(D, D1, D2)** *Fold change from basal* ($HORMONE\ ratio = max[HORMONE_{stim} / [HORMONE_{basal}]$: Expresses the hormone response as a ratio of maximal secretion to basal secretion.

Together, these analyses reveal that T2D islets display significantly higher basal hormone levels and reduced relative and fold responses to glucose, highlighting impaired stimulus-secretion coupling in both insulin- and somatostatin-secreting cells. Statistical analysis: Unpaired t-tests

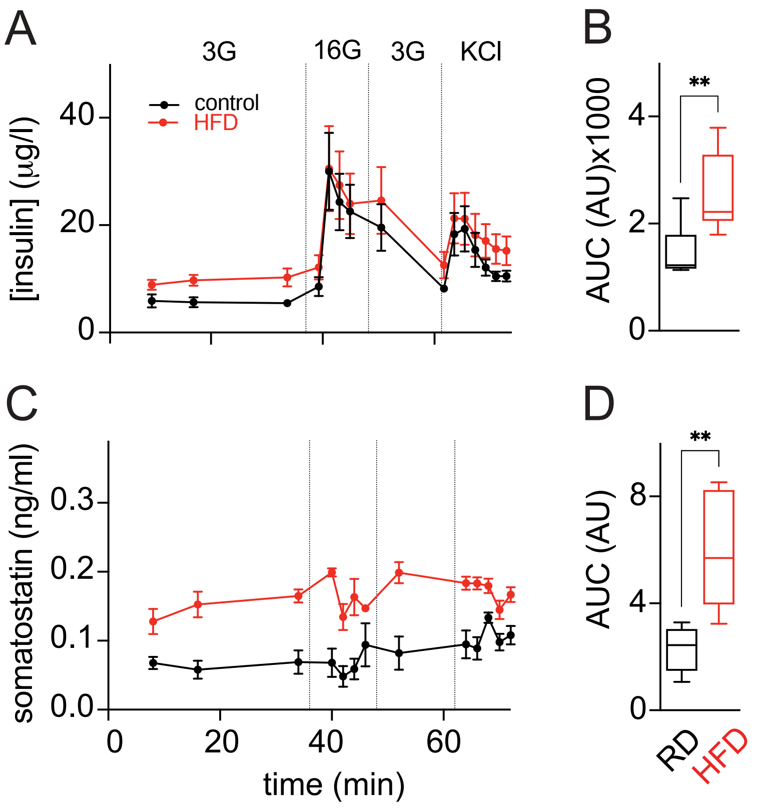


**Figure S2. Freshly Isolated Islets from High-Fat-Diet (HFD)–Fed Mice Exhibit Elevated Basal Insulin and Somatostatin Secretion with Impaired Glucose Responsiveness**. Islets were isolated from male C57BL/6J mice (Harlan Laboratories) fed either a high-fat diet (HFD) for 15 days or a regular chow diet (RD, control).

**(A, C)** Dynamic perifusion profiles of insulin (A) and somatostatin (C) secretion during sequential exposure to 3 mM and 16 mM glucose. HFD islets displayed elevated basal hormone secretion under low-glucose conditions (3 mM) and a blunted response to high glucose (16 mM) compared with RD controls.

**(B, D)** Quantification of total hormone release by area-under-the-curve (AUC) analysis across glucose transitions (3 → 16 → 3 mM) revealed significantly higher absolute secretion of insulin (B) and somatostatin (D) in HFD islets.

Data represent $n = 3$ independent perifusion experiments, each performed using pooled islets from two mice per group. Statistical significance was assessed by two-way ANOVA for dynamic secretion traces (A, C) and unpaired $t$-test (or one-way ANOVA, as appropriate) for AUC analyses (B, D). Values represent mean ± SEM. $P < 0.05$ was considered significant.

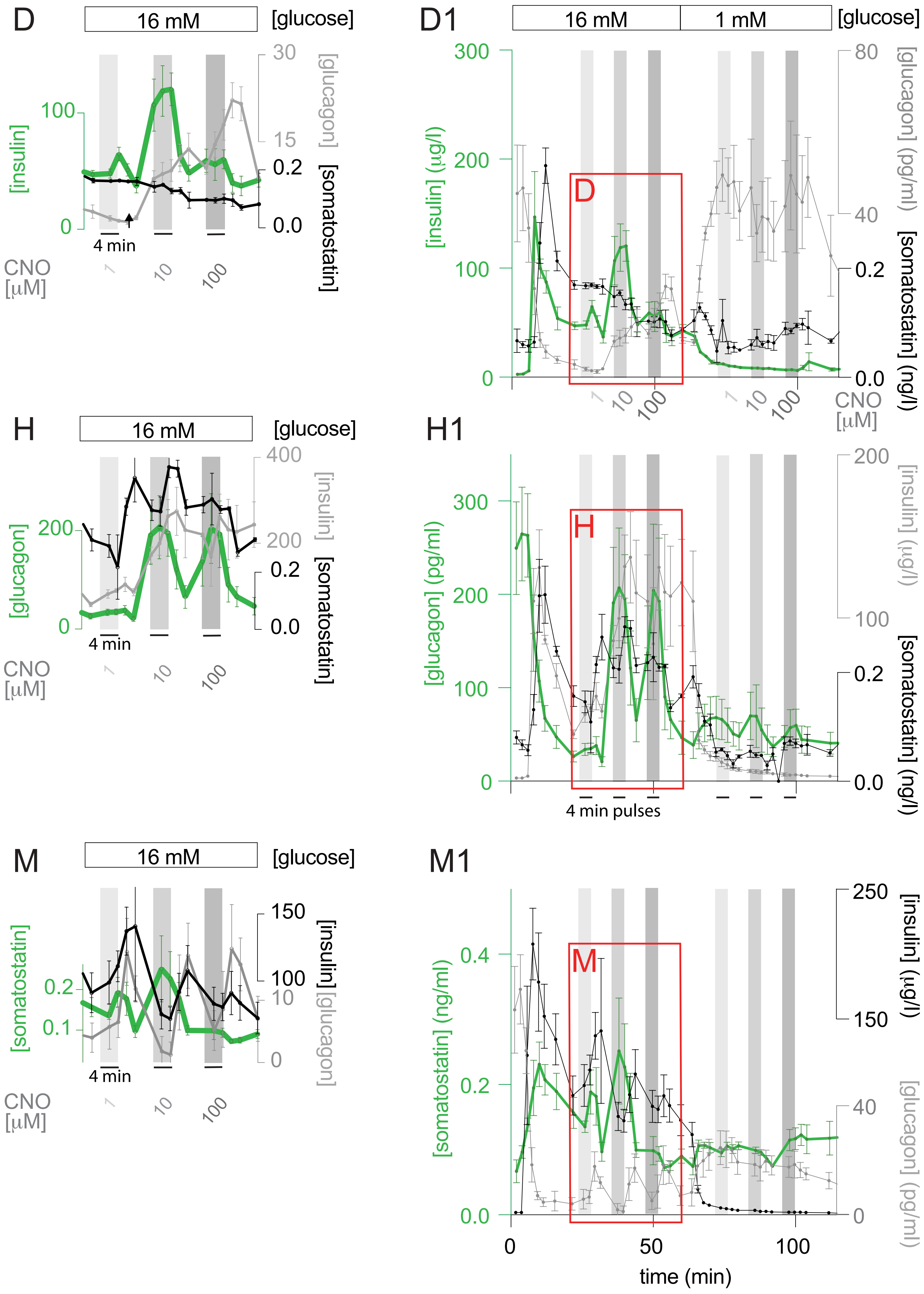

D
16 mM
[glucose]
[insulin]
[glucagon]
[somatostatin]
4 min
CNO [µM]
1
10
100
D1
16 mM
1 mM
[insulin] (µg/l)
[glucagon] (pg/ml)
[somatostatin] (ng/l)
H
H1
[glucagon] (pg/ml)
[insulin] (µg/l)
4 min pulses
M
M1
[somatostatin] (ng/ml)
time (min)

**Figure S3. Cell-Specific DREADD-Gq Activation Drives Dose-Dependent Idiotypic Secretion and Secondary Modulation of Heterotypic Hormones** (Related to Figure 2)
Chemogenetic activation of DREADD-Gq in pancreatic α-, β-, or δ-cells triggers robust, dose-dependent secretion of their idiotypic hormones—glucagon, insulin, and somatostatin—upon clozapine-N-oxide (CNO) stimulation, and modulates heterotypic hormone release during post-stimulation phases (enlargements of Figure 2D, 2H, and 2M).

**(D, H, M)** Mean ± SEM hormone secretion profiles (n = 3 donors per group) during perifusion with 16 mM glucose, illustrating idiotypic responses (green) and concurrent heterotypic dynamics (gray, black). Data represent magnified views of D1, H1, and M1.

**(D1, H1, M1)** Full perifusion traces showing pre-stimulation (3 mM glucose; −60 to 0 min), stimulation (16 mM glucose; from 0 min), and CNO pulse delivery. Curves represent group means ± SEM for each hormone measured.

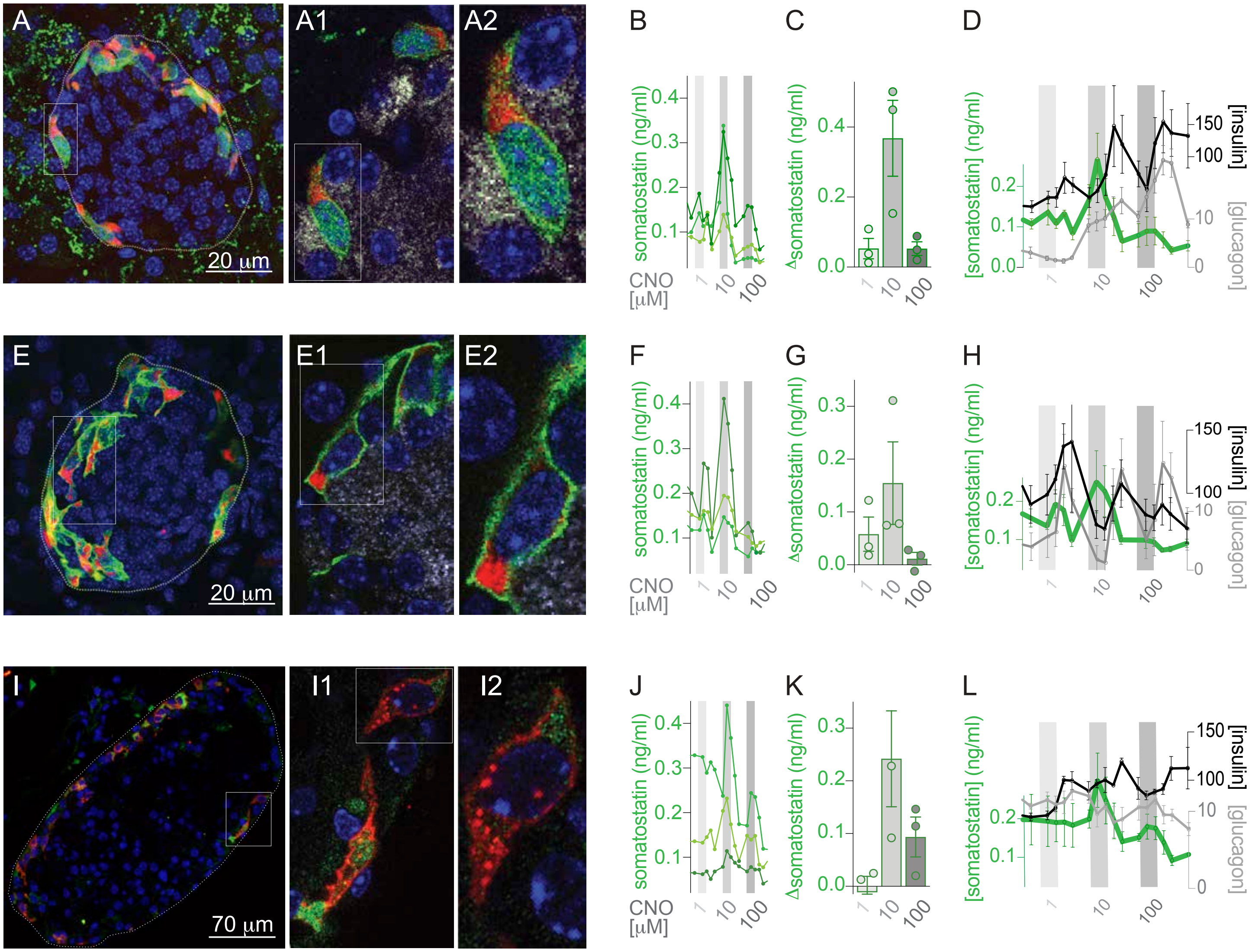

**Figure S4. Selective Expression and Functional Validation of DREADD-Gq in Pancreatic Delta Cells Across Multiple F1 Mouse Lines.** DREADD-Gq was selectively expressed in somatostatin-expressing δ-cells using three independent F1 genetic crosses: (A–D) F1[SOMATOSTATIN-Cre/DREADD-Gq–citrine-HA/GCaMP3], (E–H) F1[SOMATOSTATIN-Cre/DREADD-Gq–citrine-HA], and (I–L) F1[SOMATOSTATIN-Cre/DREADD-Gq–cherry].

**(A, E, I)** Immunofluorescence imaging of pancreatic sections shows specific expression of the DREADD-Gq reporter confined to islet δ-cells.

**(B, F, J)** Traces from perifusion assays of isolated islets demonstrate acute somatostatin secretion in response to clozapine-N-oxide (CNO) sequential stimulation.

**(C, G, K)** Quantification of somatostatin secretion reveals consistent peak responses following CNO administration across all three lines.

**(D, H, L)** Rebound secretion of insulin and glucagon is observed in islets from the three lines following somatostatin release, with temporal delays relative to somatostatin peaks .

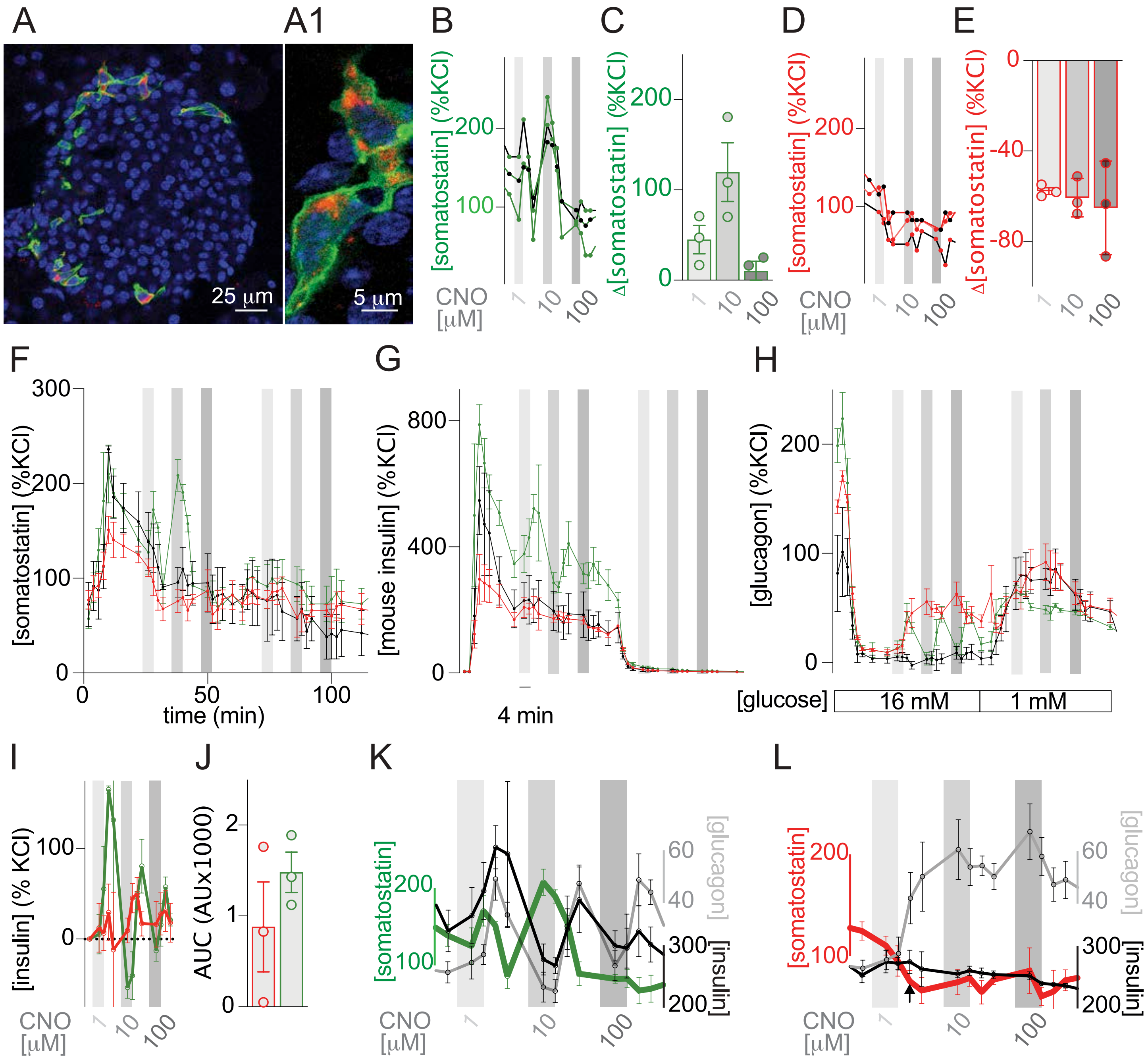


**Figure S5. KCl-Normalized Chemogenetic Modulation of δ-Cells Demonstrates Bidirectional Paracrine Regulation of Insulin and Glucagon via Somatostatin-Dependent Mechanisms.** (Related to Figure 3). All hormone secretion profiles and corresponding quantifications in this figure are normalized to the maximal secretory response to KCl, allowing comparison of relative hormone output across conditions.

Chemogenetic activation of δ-cells via DREADD-Gq enhances insulin and stimulates glucagon secretion through rebound excitatory mechanisms following acute δ-cell–mediated inhibition of α- and β-cells.

Conversely, inhibition of δ-cells using DREADD-Gi results in modest somatostatin suppression and robust disinhibition of glucagon secretion, with a comparatively smaller effect on insulin output.

**(A)** Confocal image of a pancreatic islet (scale bar, 25 μm) and **(A1)** high-resolution image of a δ-cell (scale bar, 5 μm) showing HA-tagged DREADD-Gi expression (green) localized specifically to somatostatin+ cells (red).

**(B–E)** Normalized perifusion traces and AUC quantification of somatostatin secretion in response to sequential 4-minute CNO stimulations at 1, 10, and 100 μM under 16 mM glucose in islets expressing DREADD-Gq **(B, C)** or DREADD-Gi **(D, E)**. Each trace represents an individual islet preparation (n = 3 per group), with all values expressed relative to KCl-induced secretion.

**(F–H)** Mean ± SEM hormone secretion profiles normalized to KCl response (n = 3 biological replicates per group) from islets expressing DREADD-Gq (green), DREADD-Gi (red), or no DREADD (black), depicting somatostatin **(F)**, insulin **(G)**, and glucagon **(H)** secretion under stimulatory (16 mM) and low (1 mM) glucose conditions.

**(I, J)** Estimation of excess insulin secretion following δ-cell activation or inhibition. **(I)** Subtracted traces (mean ± SEM) represent insulin output in DREADD-Gq (green) and DREADD-Gi (red) islets relative to control islets, normalized to KCl. **(J)** AUC quantification of panel I traces.

**(K, L)** Normalized secretion profiles of insulin (black) and glucagon (gray) following δ-cell activation (**K**, green) or inhibition (**L**, red), revealing reciprocal regulation of hormone output through δ-cell manipulation.

These data further support the conclusion that δ-cells exert paracrine control over both β- and α-cell activity, with somatostatin off-signaling playing a critical role in rebound hormone responses.

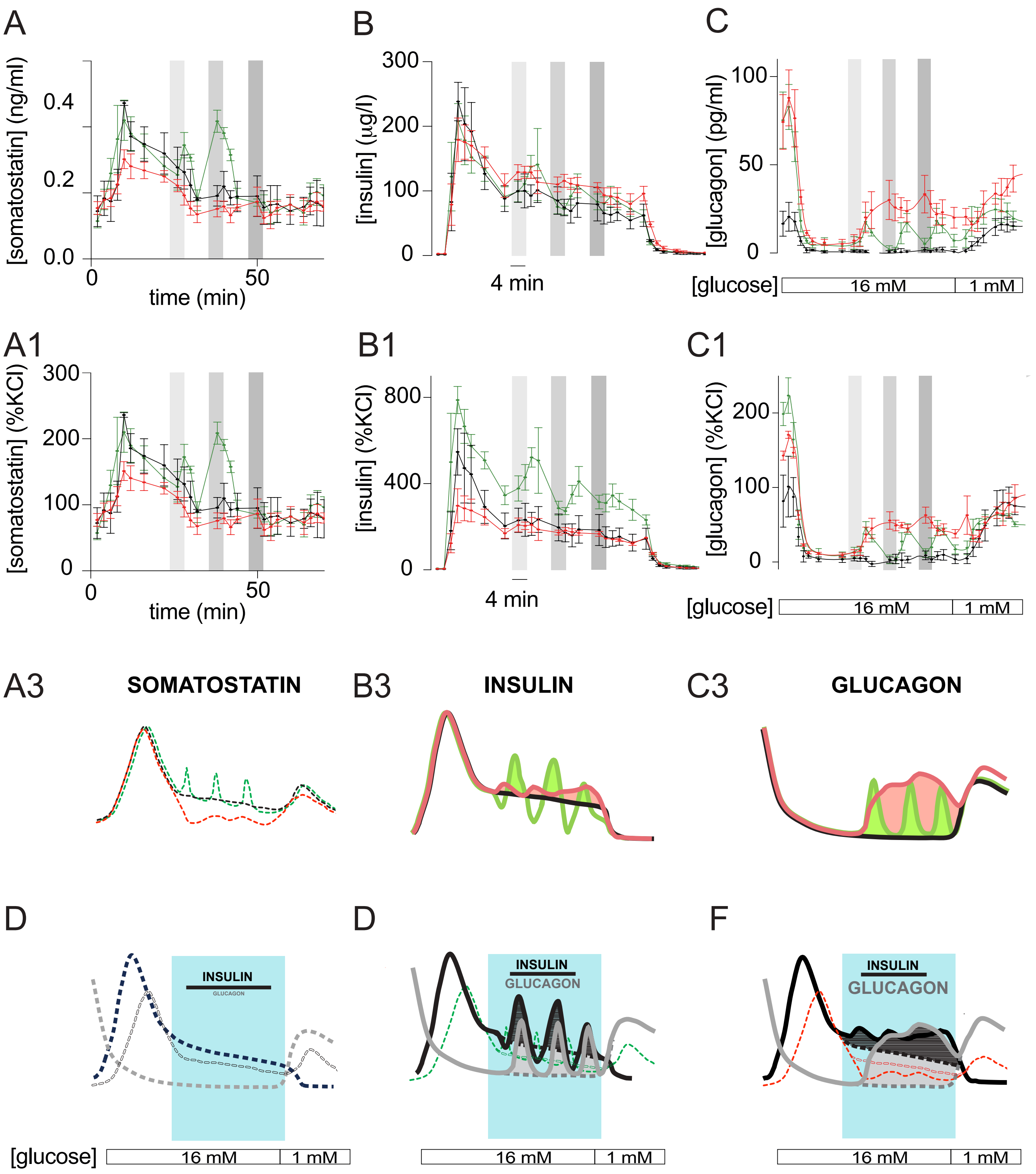
A
[somatostatin] (ng/ml)
0.4
0.2
0.0
0
50
time (min)
B
300
200
100
0
[insulin] (µg/l)
4 min
C
100
50
0
[glucagon] (pg/ml)
[glucose]
16 mM
1 mM
A1
[somatostatin] (%KCl)
300
200
100
0
0
50
time (min)
B1
800
400
0
[insulin] (%KCl)
4 min
C1
200
100
0
[glucagon] (%KCl)
[glucose]
16 mM
1 mM
A3
SOMATOSTATIN
B3
INSULIN
C3
GLUCAGON
D
INSULIN
D
INSULIN
GLUCAGON
F
INSULIN
GLUCAGON
[glucose]
16 mM
1 mM
16 mM
1 mM
16 mM
1 mM

**Figure S6. Derivation of the Conceptual Models Shown in Figure 3 from In Vitro Perifusion Experiments.** (**Related to Figure 3)**. This figure shows how the schematic models in Figure 3G were generated from *in vitro* perifusion experiments evaluating δ-cell–dependent modulation of somatostatin, insulin, and glucagon secretion under high-glucose conditions (16 mM).

**(A–C)** Representative perifusion traces of somatostatin (A), insulin (B), and glucagon (C) secretion from control (black), δ-cell–activated (green; hM3Dq + CNO), and δ-cell–inhibited (red; hM4Di + CNO) islets. Gray vertical bars indicate the timing of CNO pulses (4 min).

**(A1–C1)** Normalized hormone secretion traces from the same experiments, expressed as % of maximal KCl-evoked release, emphasize post-inhibitory rebound (PIR) patterns following each δ-cell manipulation.

**(A2–C2)** Simplified waveform reconstructions derived from the perifusion data assuming three equivalent, non-desensitizing CNO stimulations. These idealized traces illustrate hormone-specific responses: δ-cell activation evokes oscillatory rebounds (green), inhibition produces damped rebounds (red), and control islets exhibit stable low-amplitude fluctuations (black).

**(D–F)** Stylized conceptual models synthesized from the experimental waveforms summarize how δ-cell pulsatile activation (E) or inhibition (F) redistributes the relative dominance of insulin and glucagon secretion compared with control (D). Blue shaded regions represent phases of secondary insulin responses during sustained high glucose, in which δ-cell manipulations exerted their major effects. Filled areas under each curve represent net changes in hormone output relative to baseline control secretion. These models served as the templates for the schematic representations used in Figure 7 to bridge with data from systemic hormone ratios *in vivo***.**

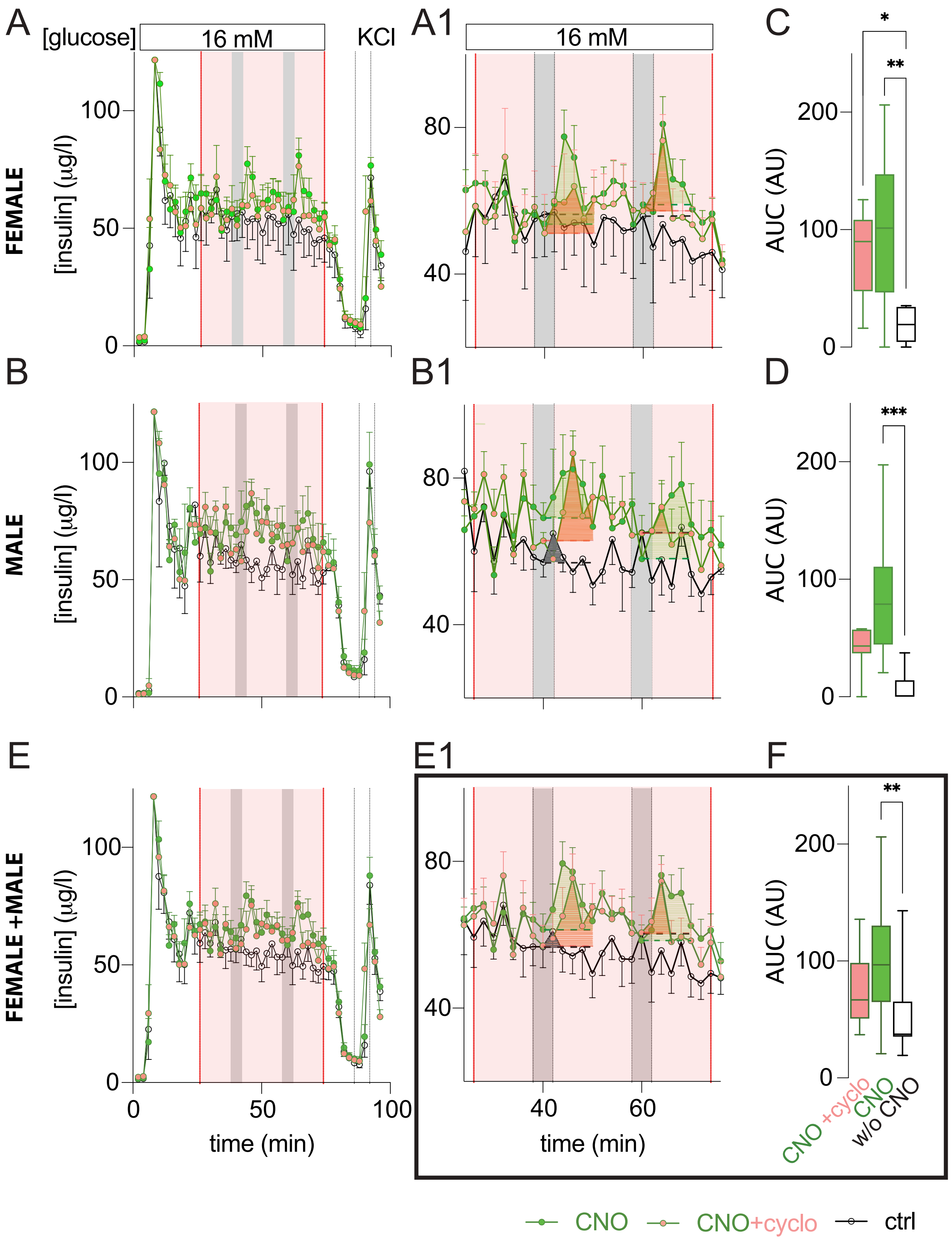


**Figure S7. Chemogenetic Activation of δ-Cells Enhances Glucose-Dependent Insulin Secretion in Both Sexes via Rebound Mechanisms Partially Mediated by Somatostatin,** Related to Figure 4 (C, D).
This figure supports that rebound insulin secretion following DREADD-Gq–mediated δ-cell activation is partially dependent on somatostatin signaling and occurs consistently in islets from both male and female donor mice.

**(A, B)** Representative perifusion traces of insulin secretion from isolated islets of female (A) and male (B) donor mice (n = 4 biological replicates per sex) stimulated with 16 mM glucose and sequential pulses of 2.5 μM CNO or sham control (black traces). δ-cell activation elicited reproducible rebound insulin secretion following each CNO pulse. Application of the somatostatin receptor antagonist cyclosomatostatin (200 nM; rose-shaded regions, applied only to DREADD-Gq–expressing islets, green trace with red-filled symbols) partially attenuated the rebound insulin response in both sexes.

**(A1, B1)** Zoomed-in views of perifusion segments highlighting the secondary insulin responses to glucose (16 mM) and CNO pulses in the absence or presence of cyclosomatostatin in female (A1) and male (B1) islets. Baseline for area-under-the-curve (AUC) quantification of the rebound insulin response was defined as insulin levels 2 min after CNO pulsing and integrated over the subsequent 10 min.

**(C, D)** Quantification of rebound insulin secretion based on AUC analysis. Bar plots depict response magnitude for female (C) and male (D) islets comparing untreated (green), cyclosomatostatin-treated (red), and non-stimulated sham control (black) groups.

**(E, E1, F)** Overlaid traces and summarized data comparing insulin secretion profiles between sexes. (E) Overlay of representative perifusion traces from both sexes. (E1, F) Quantification of rebound phases across pooled sexes confirms that somatostatin receptor antagonism partially diminishes the rebound insulin response relative to controls, indicating a partial contribution of somatostatin "off" signaling in this process. Statistical analysis: One-way ANOVA.

These results demonstrate that δ-cell activation enhances glucose-dependent insulin secretion in a sex-independent manner and provide functional evidence that somatostatin signaling modulates β-cell rebound activity under hyperglycemic conditions.

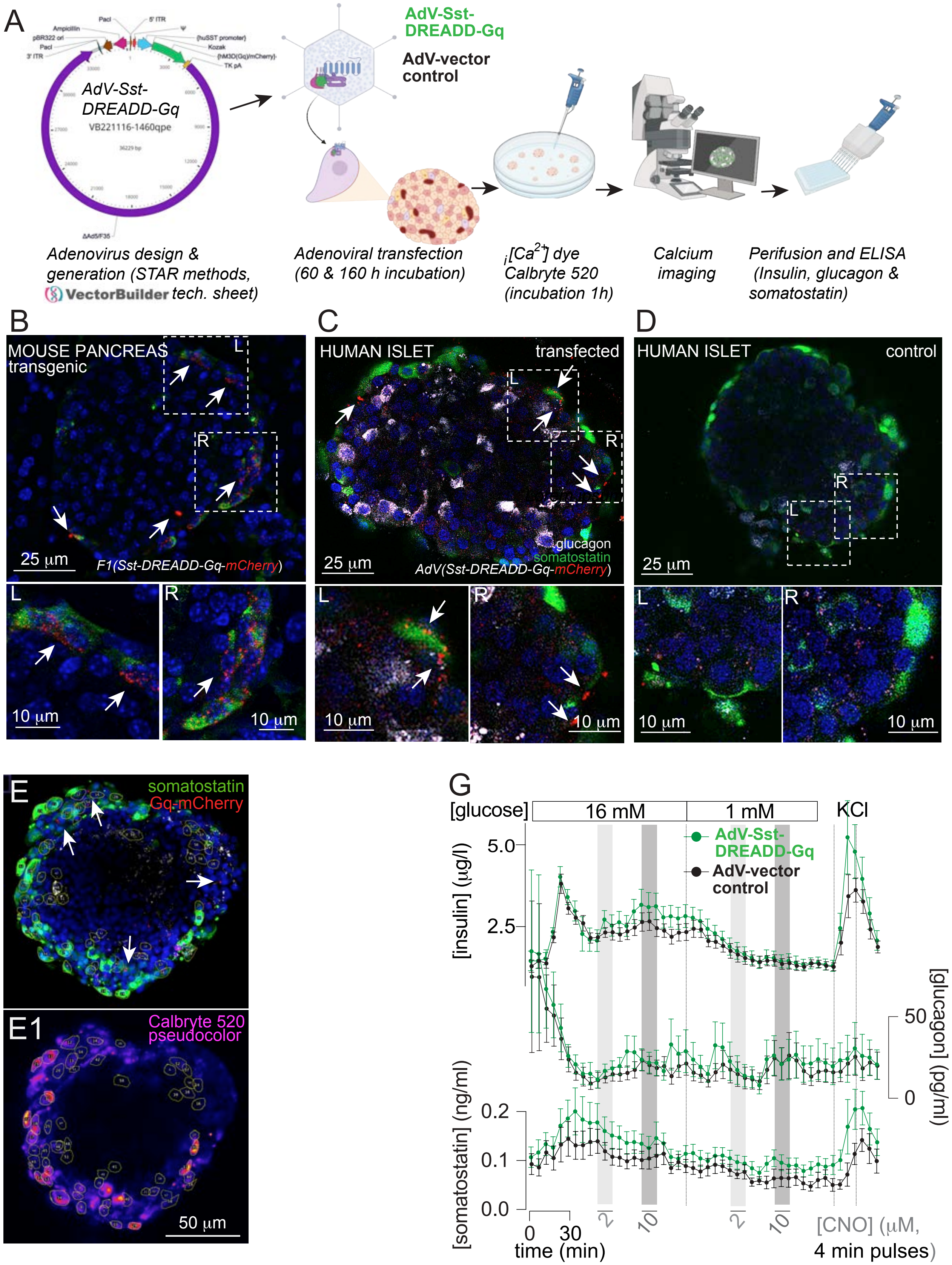

A
AdV-Sst-DREADD-Gq
VB221116-1460qpe
AdV-Sst-DREADD-Gq
AdV-vector control
Adenovirus design & generation (STAR methods, VectorBuilder tech. sheet)
Adenoviral transfection (60 & 160 h incubation)
$_i[Ca^{2+}]$ dye Calbryte 520 (incubation 1h)
Calcium imaging
Perifusion and ELISA (Insulin, glucagon & somatostatin)
B
MOUSE PANCREAS transgenic
25 μm
F1(Sst-DREADD-Gq-mCherry)
10 μm
C
HUMAN ISLET
transfected
glucagon
somatostatin
25 μm
AdV(Sst-DREADD-Gq-mCherry)
D
HUMAN ISLET
control
25 μm
E
somatostatin
Gq-mCherry
E1
Calbryte 520 pseudocolor
50 μm
G
[glucose]
16 mM
1 mM
KCl
AdV-Sst-DREADD-Gq
AdV-vector control
[insulin] (μg/l)
[glucagon] (pg/ml)
[somatostatin] (ng/ml)
time (min)
[CNO] (μM, 4 min pulses)

**Figure S8. Validation of δ-cell–targeted chemogenetic activation in human islets using AdV-SST-DREADD-Gq** (Related to Figure 4G–H). The adenoviral construct uses a human somatostatin promoter to drive δ-cell-restricted expression of hM3D(Gq)-mCherry (see VectorBuilder technical sheet for full vector design)

**(A)** Schematic of adenoviral construct design, transduction workflow, and experimental pipeline including calcium imaging and hormone secretion assays. Human islets were transduced at MOI = 50 and incubated for 60–160 h before Calbryte 520 loading, $Ca^{2+}$ imaging, and perifusion. Full vector map and sequence are provided in the Supplementary Vector Builder technical sheet.

**(B)** Validation of membrane-tagged δ-cell-specific expression of DREADD-Gq-mCherry in pancreas from transgenic mouse (F1 Sst-DREADD-Gq-mCherry) as reference system, showing co-localization of mCherry (red) with somatostatin (green). Arrows indicate the characteristic membranal dot-pattern expression of mCherry-Gq associated with δ-cells. Scale bars: 25 µm (top), 10 µm (insets).

**(C–D)** Human islet transduction with AdV-Sst-DREADD-Gq-mCherry (C) versus non-transduced control (D). mCherry (red) co-localizes with somatostatin-positive areas (green) in transfected δ-cells (arrows); glucagon (blue) marks α-cells. Unlike control islets, which show some non-specific background in red, only transfected islets displayed the characteristic membranal dot-pattern staining (arrows) observed in transgenic mice. Scale bars: 25 µm (top), 10 µm (insets).

**(E–E1)** Representative $Ca^{2+}$-imaged human islet showing somatostatin (green) and Gq-mCherry (red) expression from ICC/confocal image performed immediately after the $Ca^{2+}$-imaging procedure, using Calbryte 520 $Ca^{2+}$ pseudocolor as a broad i[$Ca^{2+}$] islet cell reporter (E1). Arrows indicate putative areas of mCherry-positive DREADD-Gq expression. Regions of interest (ROIs) encircle all responding cell areas. Scale bar: 50 µm.

**(G)** Perifusion of AdV-Sst-DREADD-Gq–transduced human islets. Islets were equilibrated at 3 mM glucose and then subjected to sequential 4-min CNO pulses (2 µM followed by 10 µM) at both 16 mM and 1 mM glucose. Experiments concluded with KCl depolarization to assess maximal secretory capacity. A non-significant trend for PIR-like rebound in insulin and glucagon secretion was observed after CNO withdrawal (n = 9 islets from 3 independent non-diabetic donors). Data are mean ± SEM.

These data suggest functionally relevant δ-cell–targeted activation and PIR using AdV-SST-DREADD-Gq in human islets.

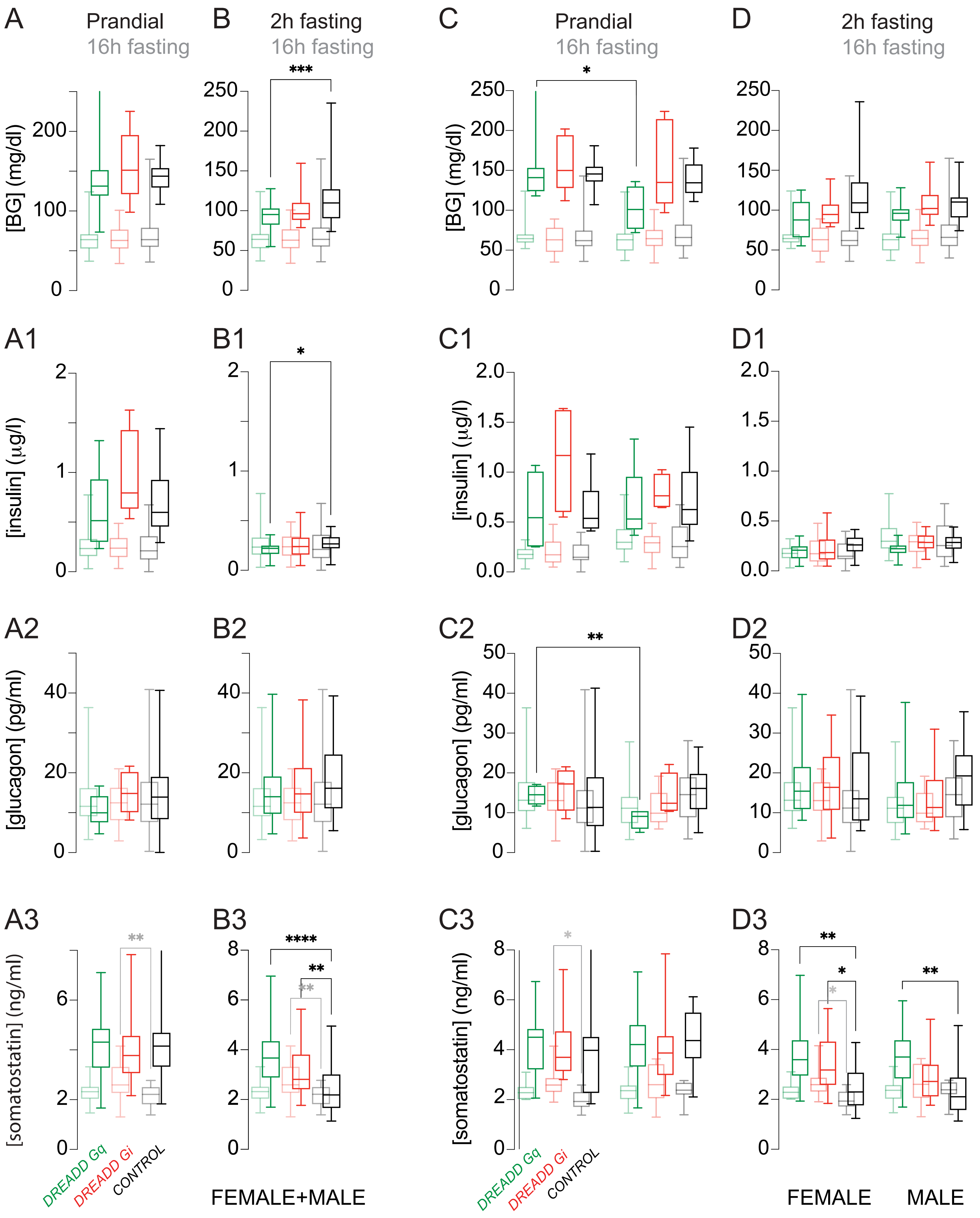

A
Prandial
16h fasting
B
2h fasting
16h fasting
C
Prandial
16h fasting
D
2h fasting
16h fasting
[BG] (mg/dl)
A1
B1
C1
D1
[insulin] (µg/l)
A2
B2
C2
D2
[glucagon] (pg/ml)
A3
B3
C3
D3
[somatostatin] (ng/ml)
DREADD Gq
DREADD Gi
CONTROL
FEMALE+MALE
FEMALE
MALE

**Figure S9. Glycemia and plasma hormone profiles across feeding states in mice engrafted with islets expressing activator-, inhibitor-, or no-DREADD in δ-cells** (Related to Figure 6B–E).
Extended analysis of non-fasting (prandial), long-fasting (16 h), and short-fasting (2 h) glycemia and circulating hormone levels in female NUDE mice transplanted with islets expressing activating DREADD-Gq (green), inhibitory DREADD-Gi (red), or no DREADD (black) in somatostatin-expressing δ-cells.

**(A–B)** Box-and-whisker plots of consolidated glycemia and plasma hormone levels across all recipients, regardless of donor sex. Comparisons are shown for prandial vs long fasting (A) and prandial vs short fasting (B) conditions.
(A1–B1) Plasma insulin; (A2–B2) plasma glucagon; (A3–B3) plasma somatostatin.

**(C–D)** Donor sex-stratified data, comparing recipient glycemia and hormone levels in mice engrafted with islets from female vs male donors. Comparisons are shown for prandial vs long fasting (C) and prandial vs short fasting (D).
(C1–D1) Plasma insulin; (C2–D2) plasma glucagon; (C3–D3) plasma somatostatin.

These data extend the findings from Figure 6B–E, confirming overall consistency in glycemia and hormone profiles across donor sexes and metabolic states. Notably, fasting somatostatin levels were elevated in mice engrafted with islets whose δ-cells expressed either DREADD variant, with the most pronounced increases observed in the inhibitory DREADD-Gi group during long fasting and in the activating DREADD-Gq group during short fasting.

### S10.1 Female nude mice transplanted with islets from FEMALE donor mice

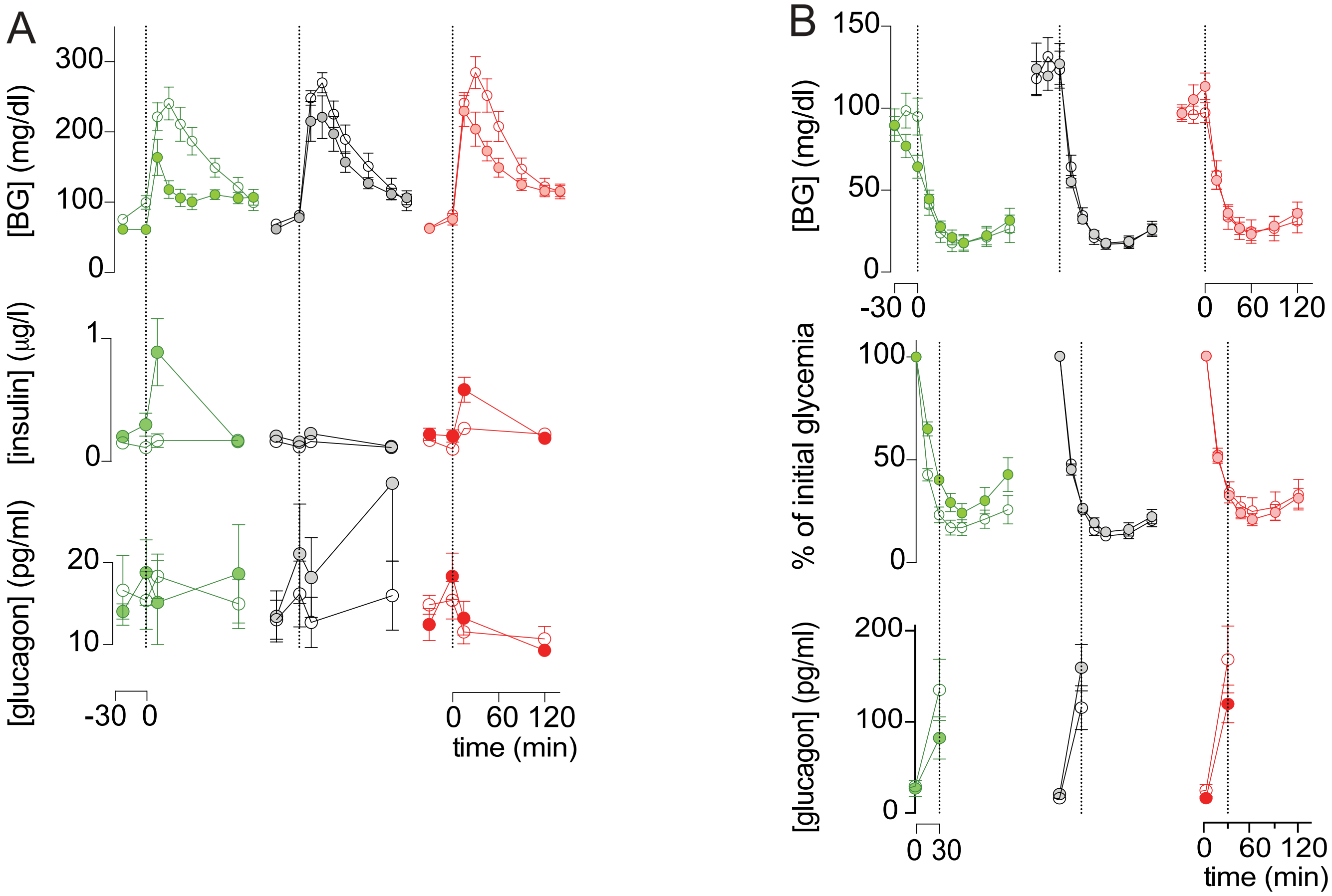


### S10.2 Female nude mice transplanted with islets from MALE donor mice

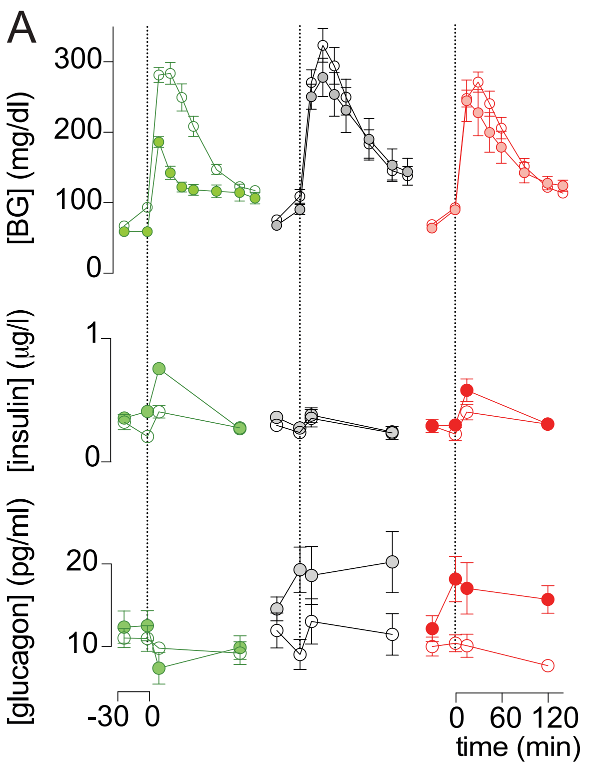


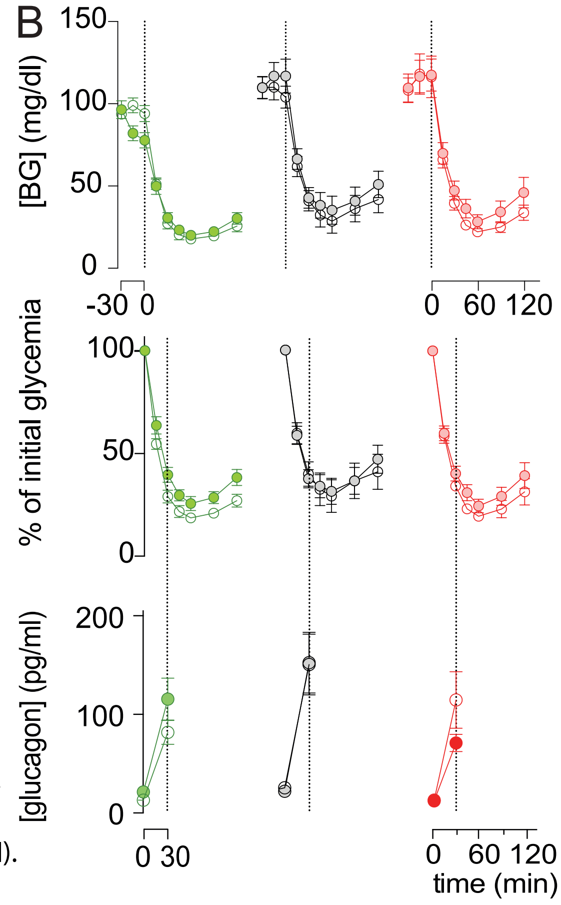


Figure S8. Acute activation, rather than inhibition, of islet δ-cells enhances insulin secretion and improves glucose handling during IPGTT and ITT across donor sexes(related to Figure 6 H–I).

**Figure S10. Acute activation, rather than inhibition, of islet δ-cells enhances insulin secretion and improves glucose handling during IPGTT and ITT across donor sexes (**Related to Figure 6H–I).

Extended analysis of glycemia and plasma hormone responses during intraperitoneal glucose tolerance tests (IPGTT; A) and insulin tolerance tests (ITT; B) in female NUDE recipient mice transplanted with islets expressing activating DREADD-Gq (green), inhibitory DREADD-Gi (red), or no DREADD (black) in somatostatin-expressing δ-cells. Data are stratified by donor sex: islets from female **(S10.1)** and male **(S10.2)** transgenic mice.

These data extend the findings from Figure 6H–I, confirming consistent glycemic responses and hormone dynamics across donor sexes. Notably, recipients of female donor islets exhibited improved glucose control during IPGTT. In both donor groups, δ-cell activation via CNO (filled green symbols) prior to glucose injection led to greater insulin release and enhanced glucose clearance compared to δ-cell inhibition (filled red symbols) or control conditions (black).

During ITT, mice engrafted with islets expressing activating DREADD-Gq showed heightened sensitivity to insulin-induced hypoglycemia under basal (non-CNO) conditions (open green symbols). However, when δ-cells were acutely activated by CNO prior to insulin injection, these same mice exhibited attenuated hypoglycemic responses (filled green symbols), indicating a shift in insulin sensitivity or counterregulatory adaptation triggered by δ-cell activation.

**Table S1. Demographic and clinical characteristics of non-diabetic donors with BMI < 26 (Lean ND) and type-2 diabetic (T2D) pancreas donors.**

| HP_ID | Age | Sex | Ethnicity | BMI | HbA1c | Cause of Death |
|---|---|---|---|---|---|---|
| HP-19342 | 64 | Male | Caucasian | 20.9 | 5.5 | Stroke |
| HP-19256 | 64 | Male | Caucasian | 25.5 | 5.4 | Stroke |
| HP-23301-01 | 64 | Male | Hawaiian | 25.5 | 5.2 | Stroke |
| HP-23262-01 | 64 | Male | Caucasian | 23.2 | 4.8 | Stroke |
| HP-21079 | 59 | Male | Caucasian | 22.6 | 5.2 | Stroke |
| HP-21079 | 59 | Male | Caucasian | 22.6 | 5.2 | Stroke |
| HP-20164 | 57 | Male | Caucasian | 24.8 | 5.1 | Stroke |
| HP-20281 | 55 | Male | Caucasian | 25.6 | 4.5 | Stroke |
| HP-20179 | 54 | Female | Hispanic | 24.6 | 5.7 | Stroke |
| HP-23278-01 | 54 | Female | Caucasian | 23.2 | 5.6 | Anoxic event |
| HP-21024 | 51 | Male | Caucasian | 20.3 | 5.2 | Stroke |
| HP-20191 | 39 | Female | Caucasian | 21.9 | 5.6 | Head trauma |
| HP-20341 | 35 | Male | Caucasian | 18.1 | 5.3 | Knife wound |
| HP-18063 | 30 | Female | Caucasian | 18 | 4.4 | Anoxia (overdose) |
| HP-21155 | 18 | Male | Hispanic | 20.4 | 5.4 | Head trauma |
| HP-19034-01T2D | 66 | Female | - | 24.9 | 5.9 | Head trauma (fall) |
| HP-19078-01T2D | 66 | Female | Caucasian | 30.3 | 6.5 | Stroke |
| HP-19171-01T2D | 66 | Female | Hispanic | 29.2 | 7.2 | Stroke |
| HP-18165-01T2D | 63 | Male | Asian | 22 | 7.3 | CVH |
| HP-21263-01T2D | 62 | Male | Caucasian | 34.6 | 6.8 | Stroke |
| HP-18320-01T2D | 61 | Male | African American | 27.4 | 7.1 | Stroke |
| HP-22193-01T2D | 60 | Female | Hispanic | 29.9 | 7.3 | Stroke |
| HP-20259-01T2D | 59 | Male | Hispanic | 25.1 | 7.3 | Head trauma |
| HP-19131-01T2D | 58 | Male | Caucasian | 32.7 | 6.7 | Stroke |
| HP-21091-01T2D | 56 | Male | Hispanic | 23.6 | 5.7 | Heart attack |
| HP-22044-01T2D | 55 | Male | Asian | 30.7 | 6.5 | Stroke |
| HP-19051-01T2D | 53 | Male | Hispanic | 30.1 | 7.8 | Head trauma (MVA) |
| HP-18243-01T2D | 51 | Male | Caucasian | 37.1 | 6.2 | Anoxic event (choking) |
| HP-21342-01T2D | 48 | Male | Hispanic | 39.2 | 7.1 | Anoxic event |
| HP-24104-01T2D | 44 | Male | Hispanic | 34.7 | 6.5 | Stroke |
| HP-21035-01T2D | 36 | Female | - | 33 | 8.3 | Stroke |
| HP-18103-01T2D | 35 | Female | Hispanic | 34 | 7.1 | Anoxia (CVA) |
| HP-18275-01T2D | 30 | Female | Hispanic | 40.1 | 6.5 | Anoxic event (asthma) |

**Table S1.** Demographic and clinical characteristics of non-diabetic donors with BMI < 26 (Lean ND) and type-2 diabetic (T2D) pancreas donors.
Individual donor data are shown for age, sex, ethnicity, BMI, HbA1c, and cause of death. The cohorts are generally well-matched for age (Lean ND: mean 49.9 ± 4.1 years, median 55.0 [Q1–Q3: 39.0–59.0]; T2D: mean 53.8 ± 2.6 years, median 57.0 [Q1–Q3: 48.8–61.8]) and sex (% male: Lean ND 69% vs. T2D 61%). Ethnicity

was more diverse in the T2D group (50% Hispanic vs. 15% in Lean ND). Stroke was the predominant cause of death in both groups (62% in Lean ND and 50% in T2D). All donors were sourced from PRODO.
Note: Detailed information on diabetes duration, treatment, severity, or degree of glycemic control was not consistently available for all T2D donors and could not be reliably obtained; this represents a limitation of the dataset.

**Table S2. Summary of Experimental Logic**

| **Experimental Arm** | **Aim** | **Manipulation** | **Readouts** |
|---|---|---|---|
| Human islet perifusion | Assess parallel insulin–somatostatin dynamics | Glucose steps | Hormone output & secretion kinetics |
| Mouse RD vs HFD islet perifusion | Test basal/stimulated hypersecretion | Diet manipulation | Hormone output |
| α/β/δ DREADD-Gq / δ DREADD-Gi characterization / intra islet cell communication | Dissect α–β–δ cell crosstalk in the islet & dissect δ-cell effects on α/β cells | CNO pulse trains / chemogenetics | PIR and rebound dynamics, co-stimulation and disinhibition surges. New Model for Paracrinicity |
| Glucose, Cyclosomatostatin Ligand (Ucn3, Ghr) | Test endogenous δ-cell-driven PIR / Physiological context | Pulsed native ligands / antagonist | Rebound insulin secretion |
| Human AdV-Sst-DREADD-Gq transduction | δ-cell specific chemogenetic activation in human islets | AdV-Sst-DREADD-Gq transduction + CNO/perifusion | Immunocytochemistry, triple-hormone secretion. Rebound dynamics. |
| Human AdV-Sst-DREADD-Gq transduction | Cellular mechanism | AdV-Sst-DREADD-Gq + CNO, Calbryte520 | Intracellular $Ca^{2+}$ transients (bulk imaging), Rebound insulin secretion |
| Intraocular graft model chemogenetic manipulation | Evaluate systemic relevance of specific δ-cell manipulation | CNO ip injected δ-cell Gq/Gi (acute) | Glycemia, plasma hormones, IPGTT/ITT |
| Intraocular graft model chemogenetic manipulation | Hormone balance | CNO in the water δ-cell Gq/Gi (chronic) | Hormone balance |
| In vivo optogenetic δ-cell activation (a-cell control) | Orthogonal validation of δ-cell-driven PIR in vivo | SST-ChR2 & GCG-ChR2 photoactivation in ACE grafts (blue light pulses) | Systemic glucose, plasma insulin |
| In vivo optogenetic δ-cell activation (a-cell control) | Targeted δ/α-cell ablation | Diphtheria-Toxin-Targeted δ/α-cell ablation | Readout specificity |
| In vivo optogenetic δ-cell activation + β-cell $Ca^{2+}$ imaging | In vivo δ-cell light activation | In vivo δ-cell activation (blue light) & b-cell-GCamP6 | Intracellular $Ca^{2+}$ transients (β-cell imaging) |
| Phenomenological modeling of δ-cell pulse–rebound dynamics | Synthesize acute data and predict chronic outcomes | Data-informed pulse-modulated SST model | Net hormone gains, INS/GCG ratio shifts |
